# Stiffness sensing fuels matrix-driven metabolic reboot for kidney repair and regeneration

**DOI:** 10.1101/2025.06.22.660927

**Authors:** Yuan Gui, Yuanyuan Wang, Wenxue Li, Jia-Jun Liu, Kelly Zheng, Jianzhong Li, Henry Wells Schaffer, Cameron Jones, Samantha Mae Mallari, Yanbao Yu, Silvia Liu, Yansheng Liu, Dong Zhou

## Abstract

Kidney repair after acute kidney injury (**AKI**) relies on a finely tuned extracellular matrix (**ECM**) that provides structural integrity and mechanical cues. As primary ECM architects, fibroblasts and pericytes rapidly mobilize to the injury site post-AKI, yet the ECM-driven repair mechanisms remain incompletely defined. Here, leveraging tissue engineering, genetic and pharmacological models, and multi-omics, we profiled the proteome landscape of decellularized kidney matrix scaffold post-AKI and highlighted microfibrillar-associated protein 2 (**Mfap2**) as a key core matrisome component primarily sourced from fibroblasts and pericytes. Mfap2 loss disrupted kidney architecture and metabolism, aggravating AKI. Global proteomics revealed that Mfap2 deficiency suppressed tubular 3-hydroxy-3-methylglutaryl-CoA synthase 2 (**Hmgcs2**) expression via estrogen receptor 2 (**Esr2**)-mediated transcriptional repression and increased protein succinylation. Phosphoproteomics and spatial transcriptomics further demonstrated a shift in mechanical signaling, with Mfap2 loss hyperactivating mitogen-activated protein kinases and upregulating large tumor suppressor kinase 1 (**Lats1**) in tubular cells without altering integrin receptor activities. In turn, Lats1 suppressed Esr2 transcription independent of its canonical Yap/Taz effectors, without affecting ubiquitin-mediated Esr2 degradation. Therapeutically, Esr2 agonists restored kidney function in Mfap2-deficient models. These findings position Mfap2 as a key regulator of ECM dynamics and mechanosignaling, linking tissue stiffness to metabolic reprogramming required for kidney repair.

## Introduction

Acute kidney injury (**AKI**) primarily affects tubular epithelial cells, long regarded as central drivers of renal regeneration(1). This reparative process unfolds within a highly dynamic microenvironment shaped not only by well-characterized biochemical and cellular responses but also by increasingly appreciated biomechanical forces, such as tissue stiffness and shear stress(2, 3). These mechanical cues are now emerging as critical regulators of early AKI repair, although the underlying mechanotransduction pathways remain incompletely defined.

Tissue stiffness, which reflects the kidney’s elasticity and resistance to deformation, is governed by the extracellular matrix (**ECM**), cellular composition, and structural architecture(4, 5). In healthy kidneys, mechanical homeostasis is maintained by a balanced ECM comprising core matrisome proteins (collagens, proteoglycans, and glycoproteins) that are structural components of the ECM that directly build the matrix scaffold thereby having a primary role in determining stiffness, and matrisome-associated proteins (e.g., MMPs and matricellular proteins) that modulate, remodel, or organize the ECM without being core structural units(6, 7). Following AKI, this equilibrium is disrupted. Fibroblasts, key producers of ECM, are among the earliest cellular responders to injury(8, 9). As the principal soft connective tissue constituents in the kidney, fibroblasts are rapidly mobilized to injury sites post-AKI(3), where they sense and respond to mechanical and biochemical cues(5, 10). In response, they synthesize context-specific ECM to scaffold tubular regeneration while concurrently transmitting mechanical signals that influence gene expression and cellular behavior through chromatin remodeling and transcriptional reprogramming(11).

Crucially, tissue stiffness is not merely a passive byproduct of injury(12). It functions as a dynamic biophysical signal that fuels a matrix-driven metabolic reboot(13), influencing the balance between repair and maladaptive fibrosis. In various organs, altered ECM composition and progressive stiffening activate mechanosensitive pathways including Yap/Taz(14–16), integrins(2, 17), and focal adhesion kinase(18, 19). Given that AKI affects approximately 22.8% of hospitalized patients globally and carries a mortality rate approaching 46% among those requiring dialysis(20–23), it is critically important to determine whether ECM-generated mechanical forces help coordinate key processes in the injury microenvironment, including fibroblast activation, epithelial plasticity, endothelium fate, immune responses, and metabolic rewiring to meet the elevated energy demands of tubular repair.

Therefore, in this study, we employed a multidisciplinary approach to systemically profile the renal core matrisome proteome landscape in AKI and to dissect the role of matrix stiffness sensing in guiding kidney repair. Our findings provide an open-access kidney matrix omics resource, offer important insights into the biomechanical regulation of renal recovery, and may inform the development of effective therapeutics reshaping the future of kidney repair.

## Results

### Characterization of proteome landscapes of decellularized kidney matrix after AKI

During AKI repair, injured tubular epithelial cells experience not only biochemical and cellular changes but also mechanical cues from the surrounding ECM and neighboring cells. Therefore, appropriate ECM stiffness is essential to support tubular cell regeneration (Figure 1A). To investigate how stiffness influences tubular cell proliferation, we cultured normal rat kidney proximal tubular epithelial cells (NRK-52E) on collagen-coated silicone gel ranging from 0.5 to 64 kPa, spanning the elastic moduli of healthy and fibrotic kidneys. Western blot assay demonstrated cell proliferative markers including cyclin B1 and D1 and their associated kinases Cyclin-dependent kinase 2 (Cdk2) and Cyclin-dependent kinase 6 (Cdk6) peaked at 2.0kPa and then were repressed along with stiffness increase (Figure 1B).

**Figure 1:**
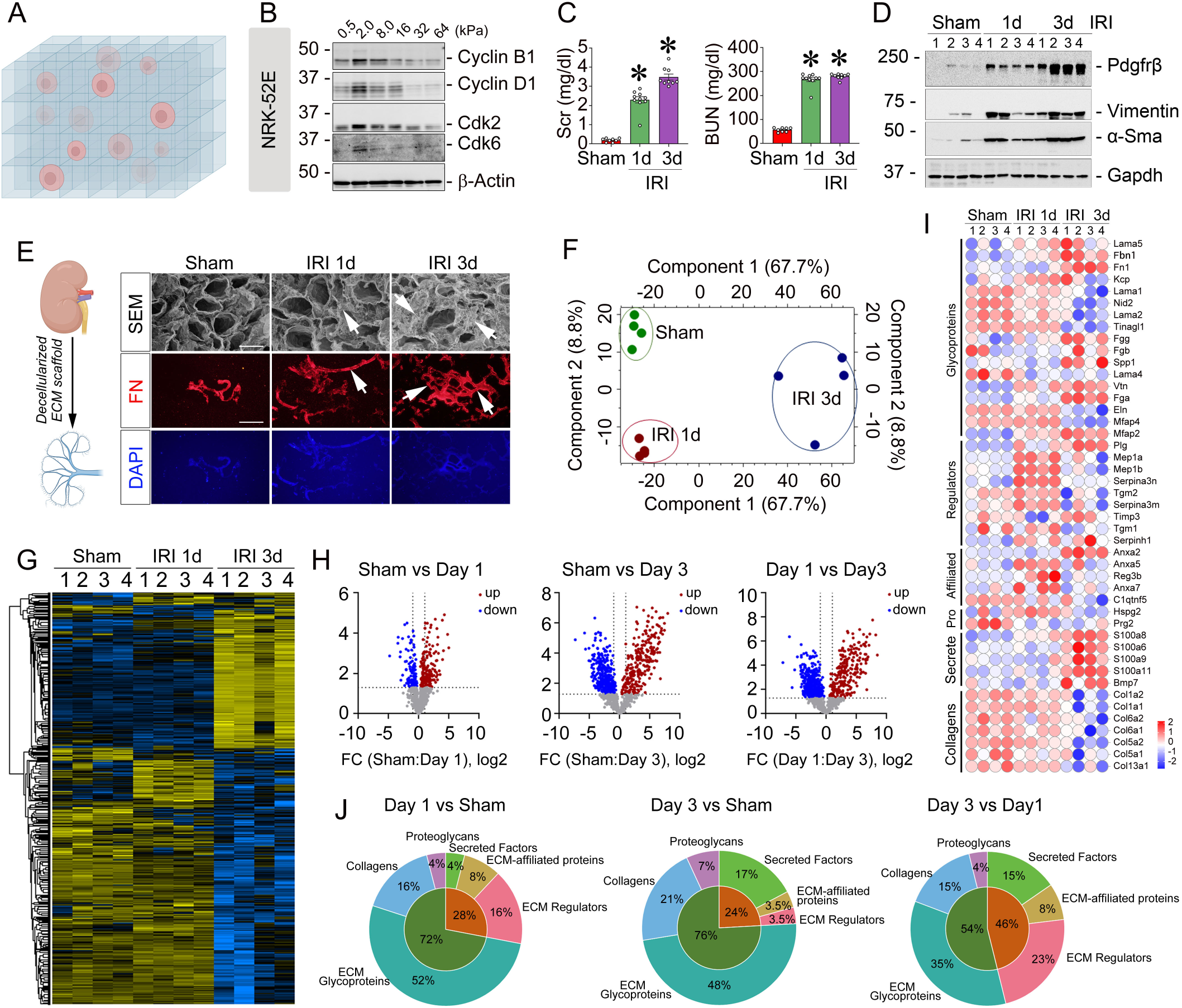
Proteomics profiles the landscape of decellularized kidney matrix scaffolds (DKS) after AKI. (A) Schematic illustration depicting matrix scaffolds within the renal microenvironment. (B) Western blot analysis of cell cycle regulators including Cyclin B1, Cyclin D1, Cdk2, and Cdk6 in normal rat kidney proximal tubular cells cultured on collagen-coated silicone gels of varying stiffness (0.5–64 kPa). (C) Serum creatinine (Scr) and blood urea nitrogen (BUN) levels in sham and IRI mice at day 0, day 1, and day 3 (n = 8-10). (D) Western blots for Pdgfrβ, Vimentin, and α-Sma proteins in sham and ischemic kidneys. Numbers denote individual animals in each group. (E) Representative transmission electron micrographs and fibronectin (FN) immunofluorescence staining of DKS from sham and ischemic kidneys. Arrows indicate positive staining. Scale bar, 50 µm. (F, G) Principal component analysis plot (F) and heatmap (G) displaying differential proteins among sham, day 1, and day 3 post-IRI groups. (H) Volcano plots of pairwise comparison between kidney proteomes of day 1 or day 3 post-IRI versus sham. Differentially expressed proteins were identified using FDR = 0.05. (I) Heatmap showing differentially expressed matrix proteins. (J) Pie chart showing the distribution of significantly altered ECM proteins between sham and IRI (day 1 and day 3) groups. * *P* < 0.05 versus sham control. Dots indicate individual animals in a given group. Graphs are presented as means ± SEM. Differences among groups were analyzed using unpaired t-tests or one-way ANOVA followed by the Student-Newman-Keuls test. IRI, ischemia-reperfusion injury; Cdk2, cyclin-dependent kinase 2; Cdk6, cyclin-dependent kinase 6; Pdgfrβ, platelet-derived growth factor receptor β; α-SMA, α-smooth muscle actin; FDR, false discovery rate.

As ECM stiffness is largely determined by core matrisome proteins, we next sought to profile the ECM proteome in the kidney. Tissue engineering strategies employing decellularized extracellular matrix provide structural and biochemical cues that recapitulate the native microenvironment(24, 25). To this end, first, we established ischemic AKI mouse models at various time points, confirming injury through elevated serum creatinine (Scr) and blood urea nitrogen (BUN) levels and increased morphological injury at day 1 and 3 (Figure 3C and Supplementary Figure S1A). Western blotting revealed increased expression of cell death markers Bax, Bad, phosphor-mixed lineage kinase domain-like (p-MLKL), as well as fibroblast and pericyte markers including platelet-derived growth factor receptor β (Pdgfrβ), vimentin, and α-smooth muscle actin (α-SMA) in the injured kidneys (Supplementary Figure S1B; Figure 1D). Then, we isolated the decellularized kidney matrix scaffold (DKS) from AKI kidneys using our established protocol(2, 26). Electron microscopy demonstrated well-preserved microarchitecture, including open tubular and vascular-like lumens surrounded by a porous, fibrous ECM, critical features for recellularization and tissue regeneration. Immunofluorescence staining confirmed enhanced ECM signal marked by fibronectin with DAPI counterstaining verifying completely removal of cellular components (Figure 1E).

To characterize the proteome-wide alternations of DKS post-IRI at day 0, 1 and 3, we performed quantitative proteomics, which led to quantitation of a total of 1,536 proteins. The reproducibility among the biological replicates was generally high, averaging over 0.9 of Pearson correlation for each experimental group (Supplementary Figure S1, C and D). An unsupervised hierarchical clustering and a principal component analysis (PCA) of the global proteome profiles clearly separate the three groups (Supplementary Figure S1E; Figure 1F), suggesting quite distinct proteomic reprogramming of the DKS under different durations of injury. When we required the proteins to be quantified in at least four out of five biological replicates, the total number of proteins reduced to 1,030, which were then utilized for further downstream analysis. A multi-sample test (ANOVA) revealed 861 significant proteins (Permutation FDR 0.05) across three time points (Figure 1G), and distinct expression patterns shown in volcano plots (Figure 1H). Focusing on ECM-related components, we filtered and annotated core matrisome and ECM-associated proteins, including structural components, regulators, and secreted factors. Heatmap clustering revealed 45 differentially expressed ECM proteins (Figure 1I). Pie chart analysis showed the core matrisome proteins comprised 72% of ECM content on day 1, increasing to 76% by day 3. While glycoprotein levels decreased by 4%, collagen content rose by 5% (Figure 1J), suggesting progressive stiffening of the kidney matrix during AKI repair. Thus, timely and well-regulated ECM remodeling is essential to foster a pro-regenerative microenvironment after injury.

### Microfibrillar-associated protein 2 (Mfap2) emerges as a key core matrisome component in the mechanical microenvironment after AKI

To further dissect the specific ECM components driving mechanical changes during AKI repair, we performed a detailed analysis of the differential ECM proteins identified in the DKS. Gene Ontology (GO) enrichment analysis across molecular function, cellular component, and biological process categories revealed that these ECM proteins were mainly associated with structural molecule activity and cell adhesion (Figure 2A). Venn diagram and heatmap showed that only six ECM proteins were substantially dysregulated at all three time points (Figure 2, B-C). Among them, Mfap2 emerged as the most highly upregulated core matrisome component in the DKS (Figure 2C). Through analyzing a public human multi-organ single-cell transcriptomic database(27), we showed minimal Mfap2 expression in healthy kidneys and livers (Figure 2, D and E). In contrast, Mfap2 expression markedly increased in murine kidneys after IRI or cisplatin-induced AKI (Figure 2, F and G). Further analyzing a separate single-nucleus RNA-seq dataset(28), fibroblasts and pericytes were identified as the predominant Mfap2-expressiong cell types after ischemic injury (Figure 2H). This cellular localization was validated by double immunofluorescence staining with Pdgfrβ (Figure 2I) in mouse models of ischemic and cisplatin-induced AKI or by immunohistochemistry staining in human AKI biopsy specimens (Figure 2J).

**Figure 2:**
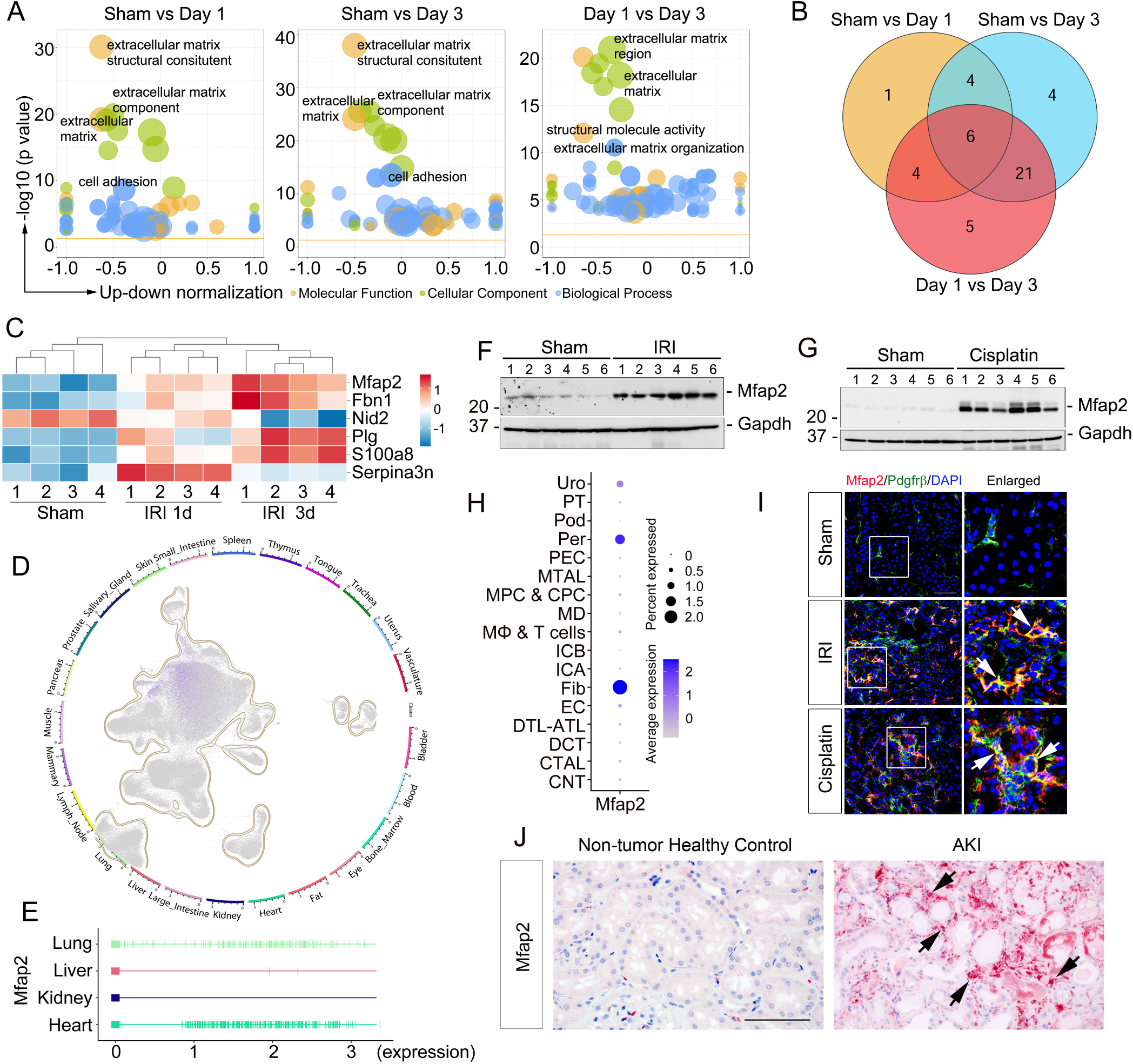
Mfap2 is a core matrisome protein upregulated in ischemic kidneys after AKI. (A) Enriched Gene Ontology (GO) terms associated with ECM protein cluster are plotted for comparison between kidney proteomes at day 1 or day 3 post-IRI versus sham. (B) Venn diagram showing the commonly expressed matrix proteins in DKS at day 1 or day 3 post-IRI versus sham. (C) Heatmap of six differentially expressed matrix proteins across sham, day 1, and day 3 after IRI. (D, E) The percentage of Mfap2-expressing cells across human organs based on a public single-cell transcriptomics atlas (GSE201333) under physiological conditions (D) and quantified by a rug plot (E). (F, G) Western blot assays of Mfap2 protein expression in the kidneys after ischemic (F) and cisplatin-induced AKI (G), respectively. Numbers denote individual animals in each group. (H) A separate public single-nucleus RNA sequencing (GSE139107) demonstrating fibroblasts and pericytes are major cellular sources of Mfap2 after IRI. (I) Double staining for Mfap2 (red) and fibroblasts/pericytes marker platelet-derived growth factor receptor β (Pdgfrβ, green) in the kidney after ischemic- and cisplatin-induced AKI. (J) Representative immunohistochemical images showed Mfap2 expression in non-tumor normal human kidney and kidney biopsy specimens from AKI patients. Arrows indicate positive staining. Scale bar, 50 µm. DKS, decellularized kidney matrix scaffolds; IRI, ischemia-reperfusion injury.

**Figure 3:**
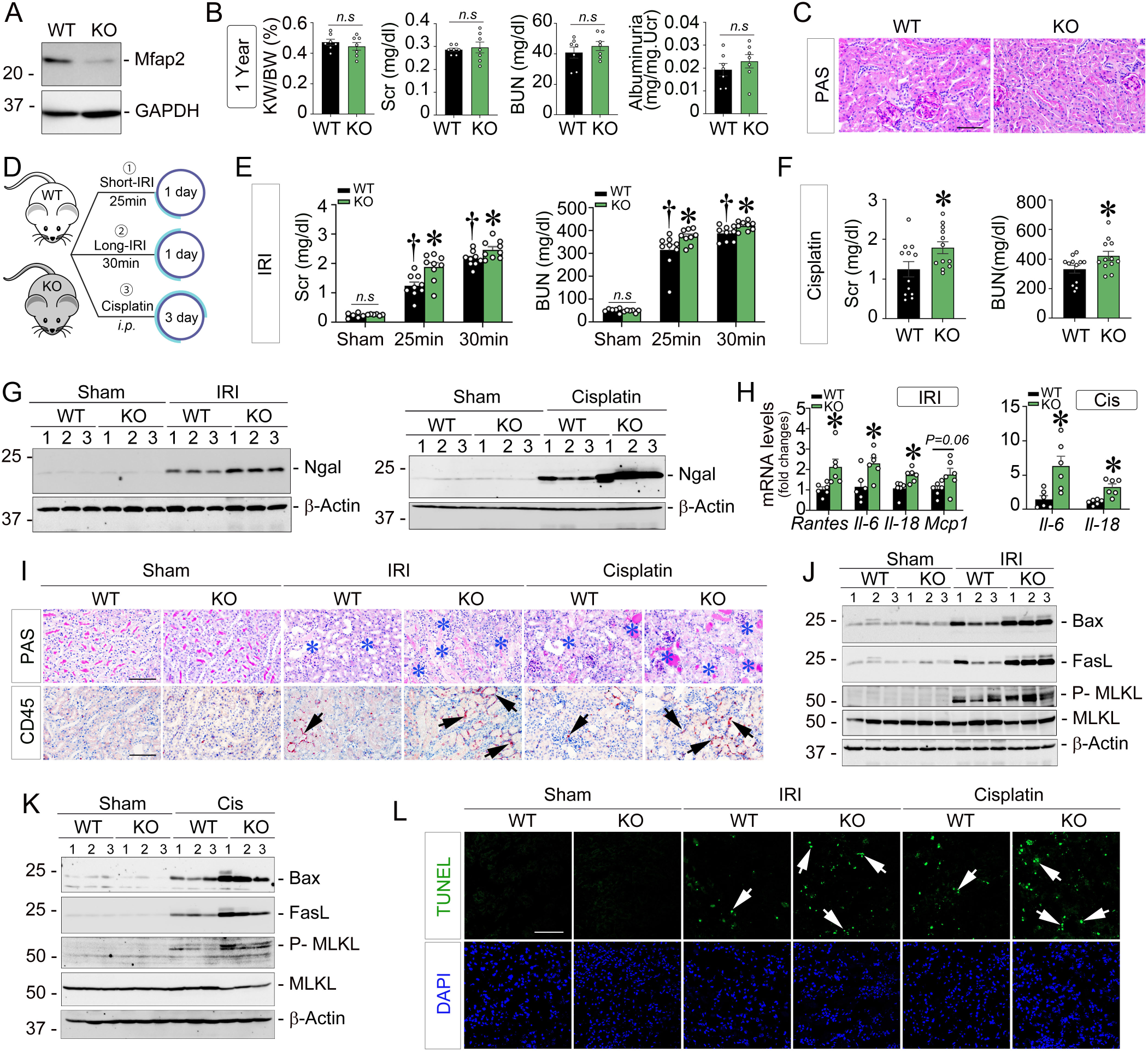
Mfap2 deficiency aggravates AKI. (A) Western blot showing Mfap2 levels in wild type (WT) and Mfap knockout (KO) kidneys. (B) Kidney weight/body weight ratio (KW/BW), serum creatinine (Scr), blood urea nitrogen (BUN), and albuminuria levels in 1-year-old WT and Mfap2 KO mice (n = 7). (C) Periodic acid–schiff (PAS) staining showing no histological differences between groups. Scale bar, 25 µm. (D) Experimental design schematic. i.p., intraperitoneal. (E, F) Scr and BUN levels 1 day after IRI (E) at 1 day or cisplatin injection (F) at 3 days (n=6-12). (G) Western blots of Ngal in the kidneys 1-day post-IRI (left) or 3 days post-cisplatin (right). (H) qRT-PCR showing *Rantes*, *Il-6*, *Il-18*, or *Mcp-1* mRNA levels 1-day post-IRI (left) or 3 days post-cisplatin (right). (n=6). (I) Representative PAS and CD45 immunohistochemical images; blue asterisks mark injured tubules; arrows indicate positive cells. Scale bar, 25 µm. (J, K) Western blots for Bax, FasL, p-MLKL, and MLKL proteins 1-day post-IRI (J) or 3 days post-cisplatin (K). (L) TUNEL staining showing apoptotic cells 1-day post-IRI or 3 days post-cisplatin. Arrows indicate apoptotic cells. DAPI is a nuclear counterstain. Scale bar, 50 µm.. † *P* < 0.05 - versus sham control. * *P* < 0.05. Dots indicate individual animals. Graphs are presented as means ± SEM. Differences among groups were analyzed using unpaired t-tests or one-way ANOVA followed by the Student-Newman-Keuls test. Rantes, regulated upon activation, normal T cell expressed and secreted; Il-6, interleukin 6; Il-18, interleukin 18; Mcp-1, monocyte chemoattractant protein-1; FasL, fas ligand; MLKL: mixed lineage kinase domain-like protein; p-MLKL: phosphor-mixed lineage kinase domain-like protein; TUNEL, terminal deoxynucleotidyl transferase dUTP nick end labeling.

### Mfap2 deficiency exacerbates AKI

To explore the functional role of Mfap2 in AKI, we generated global Mfap2 knockout (KO) mice (Figure 3A and Supplementary Figure S2A). Despite high developmental expression, Mfap2 KO mice showed no overt physiological abnormalities or renal defects at one year of age, as evident by the ratio of kidney/body weight, serum creatinine, blood urea nitrogen, and albuminuria levels (Figure 3B), as well as kidney morphological changes (Figure 3C). Then, adult Mfap2 KO mice and wild type (WT) littermates were subjected to three AKI models induced by 25- or 30-minute IRI for 1 day and cisplatin injection for 3 days (Figure 3D). Impressively, across all models, Mfap2 KO mice exhibited elevated Scr and BUN levels (Figure 3, E and F). Western blotting showed increased levels of neutrophil gelatinase-associated lipocalin (Ngal), a classic tubular injury marker, in KO kidneys (Figure 3G; quantitative data shown in Supplementary Figure S2B). Quantitative real-time PCR (qRT-PCR) analysis revealed upregulation of pro-inflammatory chemocytokines including regulated upon activation normal T expressed and secreted (*Rantes*), interleukin-6 (*Il-6*), interleukin-18 (*Il-18*), or monocyte Chemoattractant Protein-1 (*Mcp1*) in KO mice kidneys following injuries (Figure 3H). Consistently, PAS staining showed greater tubular damage in KO kidneys, with more extensive CD45+ monocyte infiltration (Figure 3I; quantitative data shown in Supplementary Figure S2C and S2D). Apoptosis- and necroptosis-related proteins including Bax, FasL, p-MLKL were also increased in KO mice after IRI (Figure 3J) and cisplatin exposure (Figure 3K). Terminal deoxynucleotidyl transferase dUTP nick end labeling (TUNEL) staining consistently showed increased apoptotic cells in KO kidneys (Figure 3L). The quantitative data are presented in Supplementary Figure S2E to S2G. In addition, Glutathione peroxidase 4 (GPX4) has no changes between WT and Mfap2 KO kidneys in all AKI models (Supplementary Figure S2H–S2K).

### Global proteomics reveals suppressed Hmgcs2-mediated ketogenesis in Mfap2-deficient kidneys post-AKI

To elucidate the mechanism underlying aggravated injury in Mfap2 KO mice, we performed data-independent acquisition (DIA)-based proteomics on ischemic kidneys from WT and KO mice(29–31). PCA demonstrated distinct clustering by genotype (Figure 4A). In total, 6,705 proteins were identified, among which 2,206 were differentially expressed, with 1,328 upregulated and 878 downregulated in KO kidneys (Figure 4B). Correlations between biological replicates within each group and the distribution of protein intensities are shown in Supplementary Figure S3A. GO analysis of biological process and cellular compartment highlighted metabolic alterations as key events, with the differentially expressed proteins predominantly localized to the mitochondrion and endoplasmic reticulum (Supplementary Figure S3B and S3C). Kyoto Encyclopedia of Genes and Genomes (KEGG) pathway enrichment analysis further confirmed that metabolic pathways are the most significantly dysregulated (Figure 3C). Among these downregulated proteins, 3-hydroxy-3-methylglutaryl-CoA synthase 2 (Hmgcs2), a key enzyme for ketogenesis in AKI(32), was markedly reduced in KO kidneys at day 1 after IRI (Figure 4D). The intensity of Hmgcs2 expression via this independent and unbiased approach were shown in Figure 4E and 4F. Then, we further validated Hmgcs2 changes in the employed AKI models. qRT-PCR analysis showed Hmgcs2 mRNA levels were substantially reduced in Mfap2 KO mice after IRI and cisplatin-induced AKI (Figure 4F), confirmed by western blot assay at protein levels (Figure 4G; quantitative data shown in Supplementary Figure S3D). Immunohistochemical staining further showed the reduction of Hmgcs2 in renal tubules in Mfap2 KO mice (Figure 4H). As Hmgcs2 catalyzes the rate-limiting step in ketone body synthesis, we then measured β-hydroxybutyrate (β-OHB) levels in ischemic kidneys. Mfap2 KO mice had substantially reduced β-OHB levels (Figure 4I). Consequently, ATP inductions were also reduced in KO mice (Figure 4J), given ketone bodies serve as alternative energy substrates that are converted into acetyl-CoA, fueling the tricarboxylic acid cycle and ultimately generating ATP.

**Figure 4:**
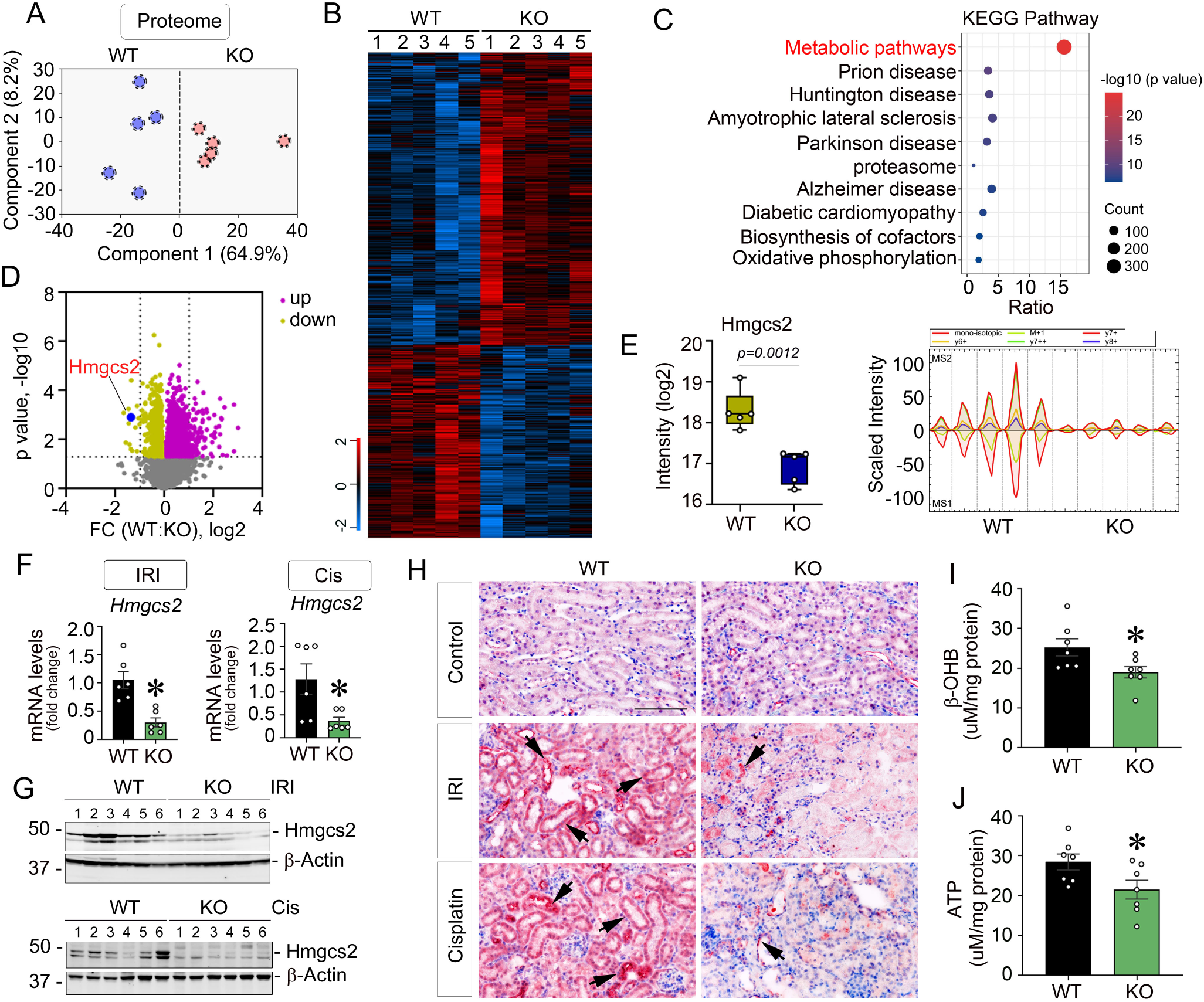
Global proteomics reveals Mfap2 deficiency suppress Hmgcs2-mediated ketogenesis in ischemic kidneys. (A) Principal component analysis of global proteomes from wild type (WT) and Mfap2 knockout (KO) kidneys 1 day after ischemic AKI. (B) Heatmap of significantly changed proteins in WT and Mfap2 KO kidneys after ischemic AKI. Two clusters of proteins with different patterns of abundance profiles are highlighted. (C) KEGG pathway enrichment analysis highlights upregulated metabolic pathways in ischemic kidneys from both groups. (D) Volcano plot showing differential proteins (Permutation FDR 0.05) between WT and Mfap2 KO kidneys after AKI. Up- and down-regulated proteins (fold-change, FC) are colored in pink and yellow, respectively. Hmgcs2 is highlighted in blue. (E) Protein intensity plot showing Hmgcs2 levels in WT and Mfap2 KO kidneys 1 day after ischemic AKI (left, n = 5). Extracted ion chromatogram (XIC) graphs from Spectronaut software for Hmgcs2 (right). MS1 XIC indicates peptide data from the first mass spectrometer, and MS2 XIC indicates peptide data from the second mass spectrometer (MS/MS). At 1 day after IRI or 3 days after cisplatin injection, (F) qPCR analysis of *Hmgcs2* mRNA in WT and Mfap2 KO kidneys (n=6). (G) Western blot showing Hmgcs2 protein expression in WT and Mfap2 KO kidneys, and numbers indicate individual animals within each group, (H) Immunohistochemical staining for Hmgcs2 in WT and Mfap2 KO kidneys. Scale bar, 50 µm. Arrows indicate positive staining. (I, J) ELISA showing β-OHB (I) and ATP (J) levels in total kidney lysates from WT and Mfap2 KO mice at 1 day after IRI (n=7). * *P* < 0.05. Dots indicate individual animals. Graphs are presented as means ± SEM. Differences among groups were analyzed using unpaired t-tests. Hmgcs2, 3-hydroxy-3-methylglutaryl-coenzyme A synthase 2; IRI, ischemia-reperfusion injury.

### Estrogen receptor 2 (Esr2) regulates Hmgcs2 activity at both transcriptional and posttranslational levels

Hmgcs2 is tightly regulated at both transcriptional and posttranslational levels(32, 33). Key transcriptional regulators include peroxisome proliferator-activated receptor α (*Pparα*), forkhead box protein A2 (*Foxa2*), and Fibroblast growth factor 21 (*Fgf21*), while Sirtuin 2 (Sirt2), Sirtuin 3 (Sirt3), and Sirtuin 5 (Sirt5) govern its posttranslational modification via acetylation or succlynation (Figure 5A). To assess changes in Mfap2 deficiency, we evaluated these factors in WT and Mfap2 KO mice kidney after IRI. Interestingly, *Pparα* and *Sirt5* mRNA levels were markedly reduced in KO kidneys, with no changes in *Foxa2*, *Fgf21*, *Sirt2*, and *Sirt3* (Figure 5B).

**Figure 5:**
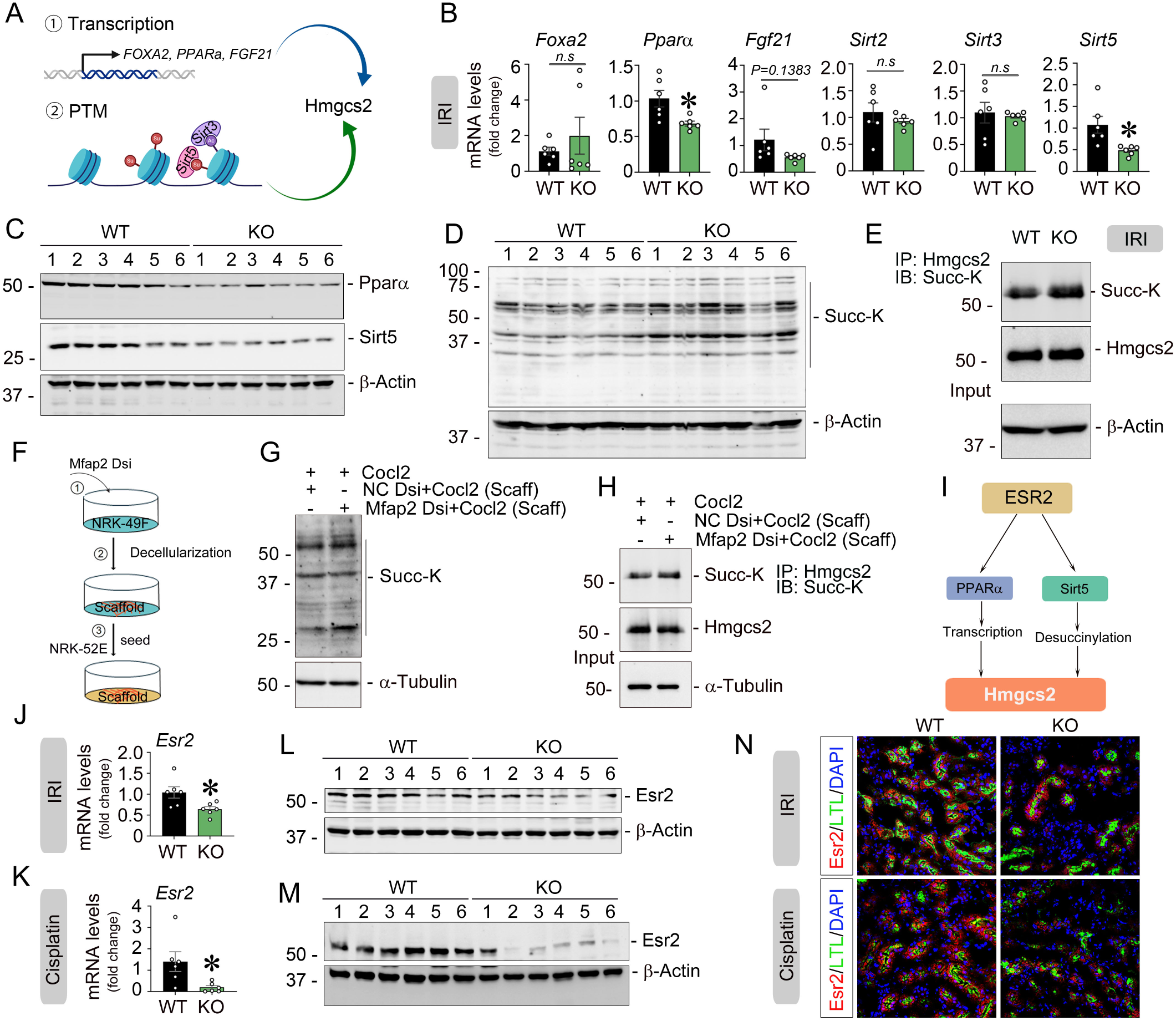
Mfap2 deletion represses Hmgcs2 transcription and lysine desuccinylation via Esr2 after AKI. (A) Schematic depicting pathways regulating Hmgcs2 expression and activity. At 1-day post-IRI, qRT-PCR of *Foxa2*, *Pparα*, *Fgf21*, *Sirt2*, *Sirt3*, and *Sirt5* mRNA in wild type (WT) and Mfap2 knockout (KO) kidneys (n=6). (B) and western blots of Pparα, Sirt5 (C), and succinylated lysine (Succ-K) (D) in WT and Mfap2 KO kidneys (n = 6). Numbers indicate individual animals within each group (C, D). (E) Immunoprecipitation of Hmgcs2 followed by succinyllysine blot showing increased lysine succinylation on Hmgcs2 in Mfap2 KO kidneys. (F) Workflow: ①NRK-49F fibroblasts were transfected with Dicer-substrate Mfap2 siRNA (Mfap2 Dsi); ② matrix scaffolds isolated; ③ NRK-52E cells seeded onto scaffolds. (G) Western blot showing increased Succ-K in NRK-52E cells on Mfap2-silenced scaffold under CoCl_2_-induced hypoxia. (H) Co-immunoprecipitation of Hmgcs2 from NRK-52E lysates showing enhanced Succ-K of Hmgcs2 in Mfap2-silenced matrix under hypoxia. (I) Schematic illustrating Esr2 regulates Hmgcs2 at transcriptional and posttranslational levels. At 1-day post-IRI or 3 days post-cisplatin, qRT-PCR showing reduced *Esr2* mRNA in Mfap2 KO kidneys versus WT (n=6) (J, K), and western blot showing Esr2 protein levels in WT and Mfap2 KO kidneys (n=6). Numbers indicate individual animals within each group (L, M). (N) Co-staining of Esr2 (red) with LTL (green) in WT and Mfap2 KO kidneys post-IRI or post-cisplatin. Scale bar, 25 µm. Arrows indicate positive cells. DAPI is used as a nuclear counterstain. * *P* < 0.05. Dots indicate individual animals in a given group. Graphs are presented as means ± SEM. Differences among groups were analyzed using unpaired t-tests. Pparα, peroxisome proliferator-activated receptor α; Fgf21, fibroblast growth factor 21; Sirt2: Sirtuin 2; Sirt3: Sirtuin 3; Sirt5: Sirtuin 5; Succ-K: succinylated Lysine; NRK-52E: normal rat kidney proximal tubular epithelial cells; Esr2: estrogen receptor 2; LTL, lotus tetragonolobus lectin.

Western blot assay confirmed the reduction of Pparα and Sirt5 in KO mice at protein levels (Figure 5C; quantitative data shown in Supplementary Figure S4A). Given our prior finding that Sirt5 modulates Hmgcs2 succinylation(32), we assessed total protein succinylation, which was elevated in KO kidneys (Figure 5D; quantitative data shown in Supplementary Figure S4B). Immunoprecipitation revealed increased Hmgcs2 succinylation in KO kidneys, suggesting reduced enzymatic activity (Figure 5E). Similarly, in cisplatin-induced AKI, Mfap2 deficiency also caused repressed expression of Pparα and Sirt 5 (Supplementary Figure S4C).

To explore why Mfap2 deficiency influences tubular Hmgcs2 levels, we used an ex vivo scaffold model. We seeded NRK-52E cells on decellularized matrix scaffold from Mfap2-silenced normal rat kidney fibroblasts (NRK-49F) (Figure 5F). Under hypoxic stress induced by CoCl2, Mfap2-silenced matrix scaffold increased protein succinylation in the seeded tubular cells (Figure 5G). Co-immunoprecipitation analysis showed similar results with in vivo study (Figure 5H).

Then, we asked whether there is a unifying upstream mechanism to determine Hmgcs2 activities. Our previous finding indicates that estrogen receptor 2 (Esr2) transcriptionally regulates Pparα or posttranslationally modifies Hmgcs2 through Sirt5 (Figure 5I)(32, 34). Therefore, we further evaluated the expression of estrogen receptor superfamily members in Mfap2 KO kidneys. Interestingly, both Esr2 mRNA and protein levels were repressed after IRI or cisplatin-induced AKI (Figure 5J to 5M; quantitative data shown in Supplementary Figure S4, D and E). Double immunostaining using Esr2 and Lotus tetragonolobus lectin (LTL) confirmed tubular localization of Esr2 and its reduction in KO kidneys (Figure 5N). However, Mfap2 deficiency did not cause changes in Esr1 expression in the diseased kidneys after AKI (Supplementary Figure S4, F and G).

### Phosphoproteomics identifies Large Tumor Suppressor Kinase 1 (Lats1) as an Esr2 regulator independent of Canonical Hippo signaling

Then, we planned to address how fibroblast/pericyte-sourced Mfap2 impacts tubular Esr2 expression after AKI. In general, cells sense mechanical cues via various mechanosensitive membrane molecules like integrins, ion channels, G protein coupled-receptors, and growth factor receptors, activating different mechanotransduction pathways(12). However, surprisingly, as a core matrisome component sourced from fibroblasts and pericytes, Mfap2 deficiency has no effect on influencing tubular integrin receptors, such as integrin α3, β1, β3, α6 (Figure 6A), as well as on activating small GTPase, including Rac1, Rac2, Cdc42, and RhoA (Figure 6B). Quantitative data are shown in Supplementary Figure S5A and S5B. However, F-actin levels were reduced and its organization disrupted in Mfap2 KO kidney, suggesting an impact on tissue stiffness after IRI (Figure 6C). Under these circumstances, to unbiasedly investigate how Mfap2 is linked to tubule-derived Esr2, we further performed DIA-based phosphoproteomics using the same set of samples for global proteomics. In total, 24,024 phospho-sites according to 4,534 proteins were identified. PCA showed clear genotype segregation (Figure 6D). Protein intensity distributions are shown in Supplementary Figure S5C, and candidate kinases that are potentially impacted are summarized in Supplementary Figure S5D. Impressively, phosphoproteomics data showed strong enrichment of mechanical signaling pathways, particularly components of the mitogen-activated protein kinase (MAPK) and Hippo signaling cascades (Figure 6E and Supplementary Figure S5E). The volcano plot showed that these phosphor-sites were upregulated in KO mice kidney after IRI (Figure 6F). Specifically, the phosphorylation of Map2k4 (Y395/S392), Map4k4 (S646/S398), and Lats1 (S612) were elevated in KO kidneys. The quantitative data were presented in Figure 6G and 6H. Interestingly, at the global level, the total proteins of Map2k4, Map4k4, and Lats1 were also elevated in KO mice after ischemic AKI (Figure 6I; quantitative data shown in Supplementary Figure S5F), whereas Map3k3, Map3k20, Map4k4, and Lats2 had no changes (Supplementary Figure S5, G and H). Most intriguingly, although the key kinase Lats1 in Hippo pathway was activated, it did not influence the canonical downstream effectors including Yap, Taz, and Tead1 (Supplementary Figure S5, I and J). Because Lats1/2 has been reported to regulate Esr1 through ubiquitination in breast cancer(35), we then investigated if it affects Esr2 upon loss of Mfap2 in AKI. To this end, we transfected Lats1 Dicer-substrate Lats1 siRNA into cultured tubular cells (Supplementary Figure S5K) and it markedly induced Esr2 expression at transcriptional levels (Figure 6J). At the protein level, knockdown of Lats1 increased Esr2 abundance and subsequently induced Hmgcs2 (Figure 6K) without affecting Esr2 ubiquitination and proteasomal degradation (Figure 6L). To further validate the transcriptional regulation between Lats1 and Esr2, we utilized Actinomycin D to repress gene transcription and found that Esr2 levels were repressed by Actinomycin D in Lats1-silenced tubular cells (Figure 6M). Western blot assay further confirmed this repression at protein levels (Figure 6N).

**Figure 6:**
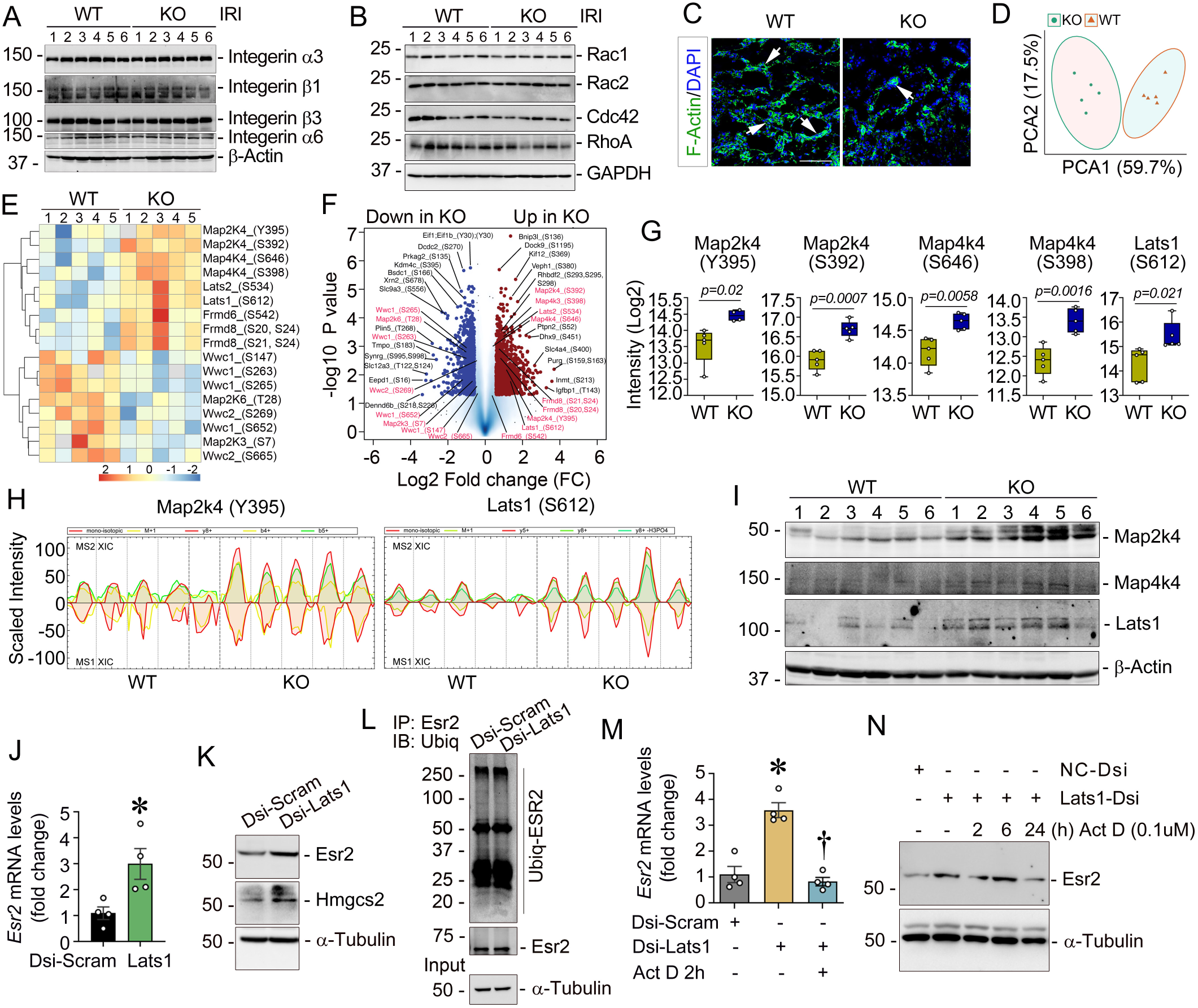
Phosphoproteomics reveals mechanotransduction activation in ischemic kidneys upon Mfap2 deletion. (A, B) Western blot showing integrins (α3, β1, β3, α6), Rac1, Rac2, Cdc42, and RhoA in wild type (WT) and Mfap2 knockout (KO) kidneys 1-day post-IRI (n = 6). Numbers indicate individual animals. (C) F-Actin in WT and KO kidneys. DAPI is used as a nuclear counterstain. Arrows indicate positive cells. Scale bar, 25 µm. (D-I) Phosphoproteomics at 1 day post-IRI: PCA plot (D), heatmap of MAPK and Hippo pathway phosphoproteins (E), volcano plot of differential phosphoproteins in Mfap2 KO kidneys (F), quantification of Map2K4 (Y395/S392), Map4K4 (S646/S398), Lats1 (S612) (G), and extracted ion chromatogram (XIC) graphs from Spectronaut software showing MS1 and MS2 phosphopeptide signals for Map2k4 (pY395, left) and Lats1 (pS612, right), representing data from the first and second mass spectrometer (MS/MS) (H). (I) Western blot of Map2k4, Map4k4, and Lats1 in WT and Mfap2 KO kidneys post-IRI (n = 6). Numbers indicate individual animals. (J-L) NRK-52E cells transfected with Dicer-substrate scramble or Lats1 siRNA (Dsi-Scramble or Dsi-Lats1) for 24 hours: qRT-PCR for *Esr2* mRNA (J), western blot of Esr2 and Hmgcs2 (K), and Esr2 ubiquitination via co-immunoprecipitation (L). (M, N) NRK-52E cells transfected with Scramble-Dsi or Lats1-Dsi were treated with Actinomycin D (Act D, 0.1µM). qPCR for *Esr2* mRNA (M) expression and Western blot of ESR2 protein levels (N). * *P* < 0.05 versus Dsi-Scramble. † *P* < 0.05 versus Dsi-Lats1. Graphs are presented as means ± SEM. Differences among groups were analyzed using unpaired t-tests or one-way ANOVA followed by the Student-Newman-Keuls test. MAPK, mitogen-activated protein kinase; Map2k4, mitogen-activated protein kinase kinase 4; Map4k4: mitogen-activated protein kinase kinase kinase kinase 4; Lats1: large tumor suppressor kinase 1; Esr2, estrogen receptor 2; NRK-52E, normal rat kidney proximal tubular epithelial cells.

### Spatial transcriptomics maps mechanical microenvironmental changes after Mfap2 loss

Having defined the mechanotransductive and signaling changes at the molecular level, we next applied 10XVisium-based spatial transcriptomics (ST) to map how Mfap2 loss reshapes the cellular and spatial landscape after AKI (Figure 7A). Overlaying ST data onto H&E-stained kidney sections (Figure 7B) revealed much higher Mfap2 expression across the cortex and medulla in WT kidneys, whereas KO kidneys exhibited a striking global loss of Mfap2 signal (Figure 7C). To decipher spatial gene expression patterns, we classified ST spots into three anatomically defined clusters: cortex, outer medulla, and inner medulla (Figure 7D). Consistent with our in vivo findings (Figure 3), the spatial profiles confirmed that the injury marker Ngal was extensively upregulated throughout the Mfap2 KO kidney, while kidney injury molecule 1 (Kim-1) remained largely unchanged in the inner medulla (Figure 7E, left and middle panels). Similar to Kim-1, Hmgcs2 levels were sharply decreased particularly in the cortex and outer medulla of Mfap2 KO kidneys (Figure 7E, right panel). Quantitative data for these markers across the three anatomical clusters are presented in Figure 7F, with the differential test between WT and KO kidneys.

**Figure 7:**
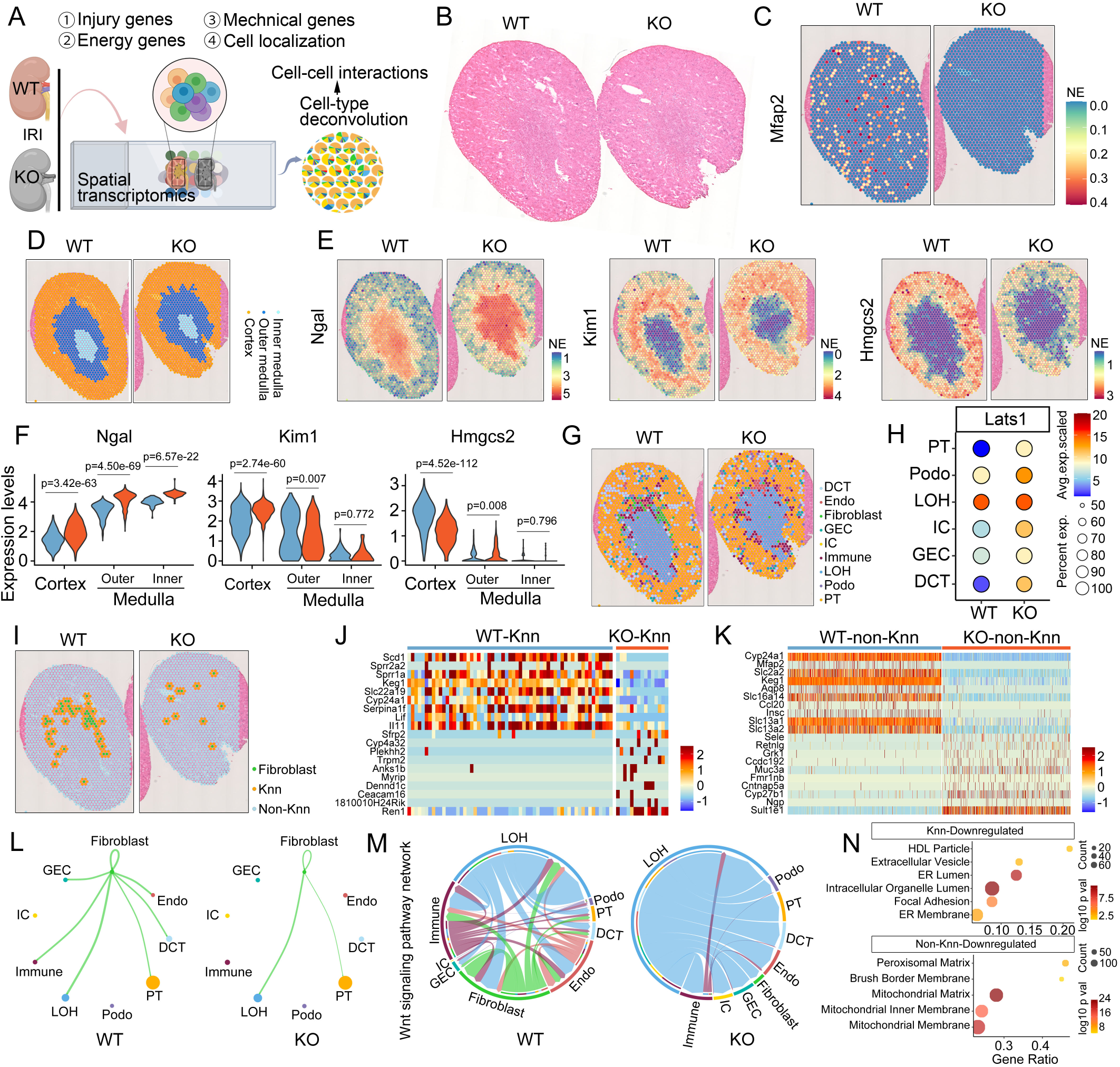
Spatial transcriptomics defines the mechanical microenvironment dynamics of injury kidneys loss of Mfap2 after AKI. (A) Overview of the experimental workflow. Spatial transcriptomics (ST) was conducted on injured kidneys from wild type (WT) and Mfap2 knockout (KO) mice to profile injury-associated genes, energy metabolism genes, and genes related to the mechanical microenvironment, cellular localization, and cell-cell interactions. (B) Hematoxylin & Eosin (H&E) staining of kidney sections from WT and KO mice post-IRI. (C) Spatial map showing Mfap2 levels in WT versus KO kidneys. (D) Spatial separation of inner medulla, outer medulla, and cortex across genotypes. (E, F) Spatial maps (E) and quantification (F) of injury markers (Ngal and Kim1) and energy metabolism marker Hmgcs2 in WT and KO kidneys. (G) Spatial deconvolution illustrating the cell types annotation per spot in WT and KO kidneys. (H) Dot plots showing Lats1 expression across nephron segments in WT and KO kidneys. (I) Spatial distribution of fibroblasts and their K-nearest-neighbor (Knn) spots clusters across genotypes. (J, K) Differentially expressed genes between WT and KO kidneys, focusing on tissue adjacent to (Knn, J) and distant from (non-Knn, K) fibroblasts. (L) Spatial cellular ligand-receptor interactions between fibroblasts and the other nephron cell types in WT and KO kidneys. (M) Alteration of FGF-mediated signaling connections across nephron and cellular compartments. (N) GO enrichment analysis under cellular compartment term showing down-regulated biological events of Knn and non-Knn regions between WT and KO mice after IRI. IRI, ischemia-reperfusion injury; PT, proximal tubule; LOH, loop of Henle; DCT, distal convoluted tubule; GEC, glomerular endothelial cell; IC, intercalated cell; Endo, endothelial cell; Immune, immune cells; FGF, fibroblast growth factor; GO, gene ontology; HDL, high-density lipoprotein; ER, endoplasmic reticulum; NE, normalized expression; Percent exp., percent expressing; Avg. exp., average expression; Val, value.

To achieve single-cell spatial resolution, we integrated our ST data with a published single-cell RNA-seq dataset from ischemic mouse kidneys (GSE180420)(36). Deconvolution analyses (Figure 7G; Supplementary Figure S6A and S6B) revealed significant shifts in cellular composition across spatial spots, with Mfap2 KO kidneys exhibiting a marked loss of proximal and distal tubular epithelial populations post-AKI. The analysis identified spatial patterns for major kidney cell types, including proximal tubules (PT), loops of Henle (LOH), distal convoluted tubules (DCT), fibroblasts, endothelial and immune cells, and intercalated cells (IC). Of note, we observed substantially increased Lats1 predominantly within PT, IC, and DCT nephron compartments in KO kidneys (Figure 7H).

Since Mfap2 is primarily expressed by fibroblasts and pericytes, we further examined spatial cell-cell communication by assessing genes associated with the six nearest neighbors (Knn) spots adjacent to fibroblasts in WT and KO kidneys (Figure 7I). The differential analysis revealed that Knn spots were enriched for mechanobiology-related genes, such as *sprr2a2*, *sprr1a*, and *keg1*, which were downregulated in Mfap2 KO kidneys (Figure 7J). In contrast, genes associated with non-Knn spots captured a broader cellular response across nephron compartments, including metabolism, ion transport, immune infiltration, and tissue polarity adjustments (Figure 7K). Most impressively, cellular ligand-receptor communication analysis revealed that loss of Mfap2 substantially weakened fibroblast-neighbor interactions, impairing the establishment of a supportive microenvironment for renal repair (Figure 7L). The Wnt signaling pathway, a well-characterized AKI ‘protector’(2, 37), was markedly suppressed in cell-cell communications within Mfap2 KO kidneys (Figure 7M). Moreover, several MAPK upstream regulators known to protect against AKI(38–41), including fibroblast growth factors (FGF), epidermal growth factors (EGF), platelet-derived growth factor (PDGF), and matrix protein tenascin, were also disrupted in Mfap2 KO kidneys (Supplementary Figure S6C). Consistently, GO analysis under cellular compartment term showed reduced activity of focal adhesion in Knn regions while mitochondrial-related pathways were downregulated in non-Knn regions in Mfap2 KO kidneys (Figure 7N), collectively exacerbating AKI severity.

### Esr2 agonist attenuates AKI

Given loss of Mfap2 reduced Esr2 and aggravated AKI (Figure 3 and 5), we next evaluated the therapeutic relevance of Esr2 signaling. Mfap2 KO mice were intraperitoneally treated with the selective Esr2 agonist, Erb-041 (1mg/kg) for two days prior to IRI (Figure 8A). Remarkably, Erb-041 substantially lowered Scr and BUN levels post-IRI (Figure 8B) and reduced expression of tubular injury marker Ngal and apoptotic markers Bax and FasL (Figure 8C; quantitative data shown in Supplementary Figure S7A). PAS staining revealed improved tubular morphology (Figure 8D), while TUNEL staining confirmed decreased apoptosis (Figure 8E). Immunohistochemical staining showed reduced infiltration of CD45+ monocytes and CD68+ macrophages (Figure 8F). Quantitative data are presented in Supplementary Figure S7, B to E. qRT-PCR analysis further demonstrated reduced mRNA levels of cytochemokines, including *Rantes*, *Il-6*, *Il-18*, and *Mcp-1* (Figure 8G). Consistently, western blot assay revealed a robust increase in Hmgcs2 protein levels (Figure 8H; quantitative data shown in Figure 8I), corroborated by enhanced tubular Hmgcs2 expression with immunohistochemistry staining (Figure 8J). This was accompanied by elevated β-OHB and ATP production (Figure 8, K and L). In vitro, Erb-041 induced Hmgcs2 in cultured tubular cells (Figure 8M). Upon hypoxic stress, Erb-041 markedly reduced Bax expression (Figure 8N). Moreover, in Staurosporine-induced apoptosis, Erb-041 markedly reduced cleaved caspase 3 levels (Figure 8O), confirming direct cytoprotective effects.

**Figure 8:**
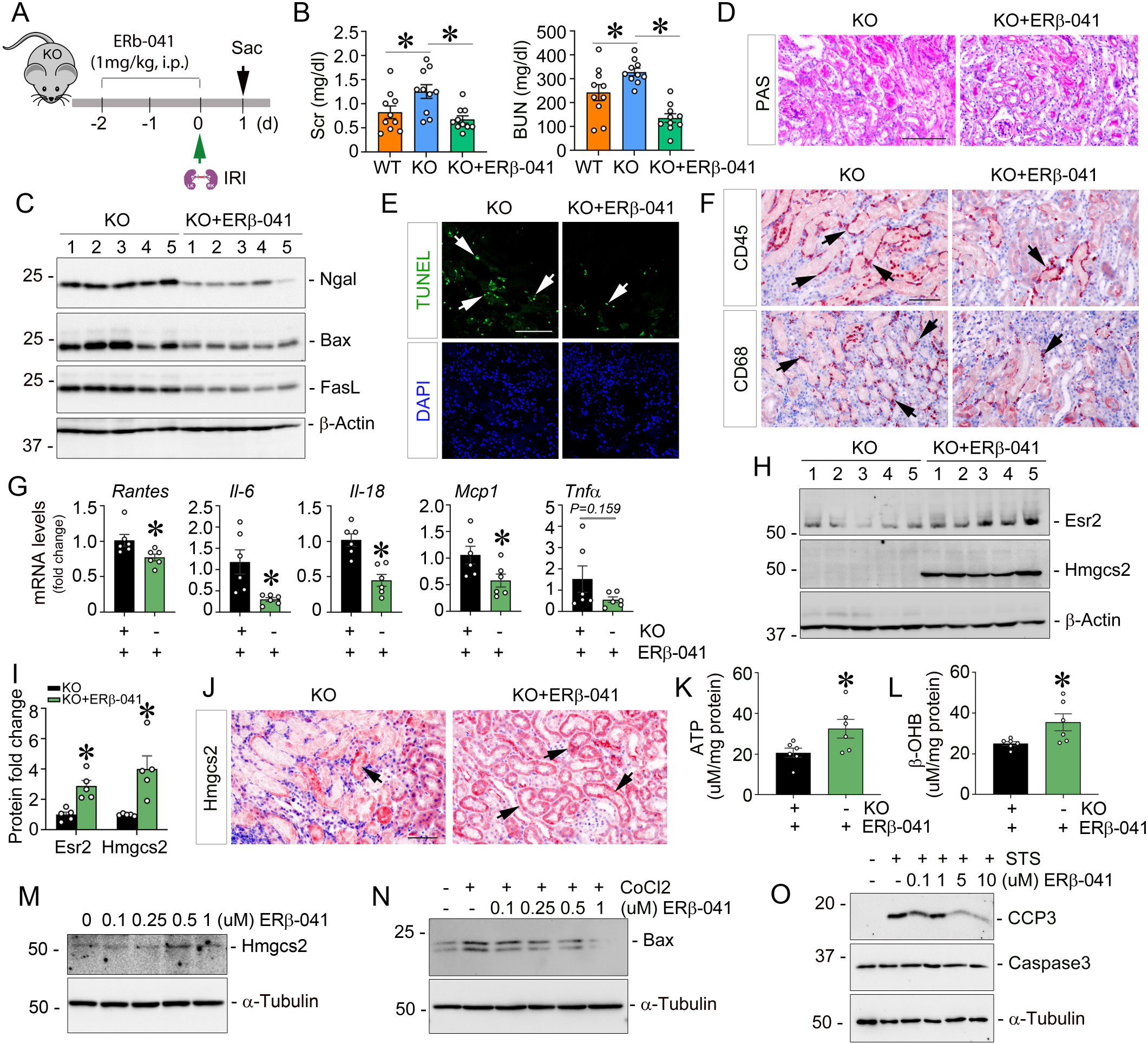
Esr2 agonist mitigates ischemic AKI. (A) Experiment design. The selective Esr2 agonist, Erb-041, was administered to Mfap2 KO mice two days prior to IRI. (B) Serum creatinine (Scr) and blood urea nitrogen (BUN) levels in WT and Mfap2 KO mice after IRI with or without Erb-041 treatment (n=10). Comparing Mfap2 KO mice with or without Erb-041 treatment, at 1 day after IRI, Western blot analysis of Ngal, Bax, and FasL in the kidneys (n = 5). Numbers indicate individual animals within each group (C); Periodic acid–schiff (PAS) staining of the kidneys. Scale bar, 25 µm (D); TUNEL and immunohistochemical staining against CD45 and CD68 in the kidneys. Scale bar, 25 µm. Arrows indicate positive cells. DAPI is a nuclear counterstain (E, F). (G) qPCR analysis of *Rantes*, *Il-6*, *Il-18*, *Mcp1*, and *Tnf-α* mRNA levels in the kidneys (n=6). (H, I) Western blot (H) and quantitative data (I) of Esr2 and Hmgcs2 proteins in the kidneys (n=5). Numbers indicate individual animals within each group. (J) Immunohistochemical staining of Hmgcs2 protein expression in the kidneys. Arrows indicate positive staining. Scale bar, 50 µm. (K, L) ELISA measurements of ATP (K) and β-OHB (L) in whole kidney tissue (n=6). (M-O) NRK-52E cells were treated with varying doses of ERb-041 showed increased Hmgcs2 expression (M), decreased Bax levels (N), and reduced cleaved caspase-3 (CCP3) (O), as determined by western blot assay. * *P* < 0.05. Dots indicate individual animals in a given group. Graphs are presented as means ± SEM. Differences among groups were analyzed using unpaired t-tests or one-way ANOVA followed by the Student-Newman-Keuls test. Esr2, estrogen receptor 2; IRI, ischemia-reperfusion injury; Ngal, neutrophil gelatinase-associated lipocalin; Rantes, regulated upon activation, normal T cell expressed and secreted; Il-6, interleukin 6; Il-18, interleukin 18; Mcp-1, monocyte chemoattractant protein-1; FasL, fas ligand; TUNEL, terminal deoxynucleotidyl transferase dUTP nick end labeling; Hmgcs2, 3-hydroxy-3-methylglutaryl-CoA synthase 2.

### Mfap2 mediates matrix-driven metabolic reboot for tubular repair in vitro and ex vivo

To validate our in vivo findings, we conducted ex vivo and in vitro studies. First, we seeded NRK-52E cells on DKS from WT and Mfap2 KO mice after IRI, respectively (Figure 9A). Upon Staurosporine treatment, tubular cells seeded on Mfap2 KO scaffolds exhibited increased apoptosis, as indicated by TUNEL staining (Figure 9B). Given the contribution of fibroblasts to matrix stiffness, we next knocked down Mfap2 in cultured fibroblasts (Supplementary Figure S8A). Under CoCl2-induced hypoxic stress, these Mfap2-deficient fibroblasts displayed disorganized F-actin structures (Figure 9C). We then removed cells and obtained fibroblast-scaffolds lacking Mfap2, confirmed by western blotting (Figure 9D). NRK-52E cells cultured on these scaffolds exhibited increased cleaved caspase-3 after −3 hours of Staurosporine stimulation (Figure 9E). Similarly, the scaffolds induced upregulation of Ngal and Bax in seeded tubular cells, both under basal conditions and in response to hypoxic stress (Figure 9F and Supplementary Figure S8B). Mechanistically, knockdown of Mfap2 elevated key mechanotransduction components, including Map2k4, Map4k4, and Lats1, resulting in Esr2 repression and impaired Hmgcs2 inductions in tubular cells (Figure 9F and Supplementary Figure S8C).

**Figure 9:**
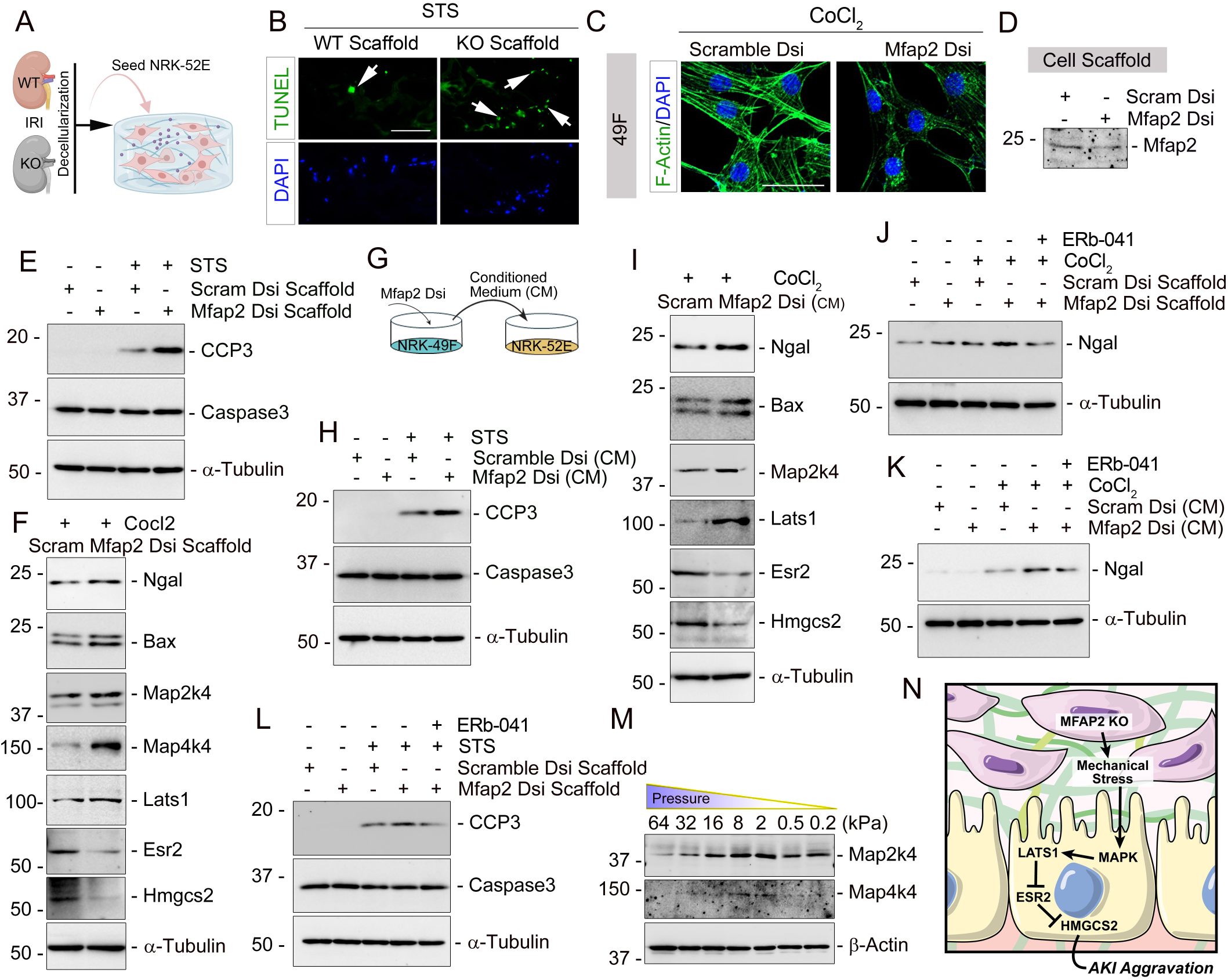
Mfap2 mediates matrix-driven metabolic reboot for tubular repair in vitro and ex vivo. (A) Schematic diagram. (B) TUNEL staining showing increased NRK-52E cells seeded on Mfap2-deficiency DKS underwent apoptosis upon Staurosporine stimulation, compared to controls. (C) NRK-49F fibroblasts were exposed to hypoxia (CoCl_2_, 400 µM) and transfected with Dicer-substrate Mfap2 siRNA (Mfap2 Dsi) for 24 hours. Immunofluorescence staining showed reduced F-Actin in Mfap2-silenced fibroblasts. Scale bar, 25 µm. (D) Western blot confirmed reduced Mfap2 protein in decellularized fibroblast-derived matrix scaffold. (E) Western blot assay of NRK-52E cells seeded on Mfap2-silenced matrix scaffold showing increased cleaved caspase-3 (CCP3). (F) Western blot showing increased Ngal, Bax, Map2k4, Map4k4, Lats1, decreased Esr2, and reduced Hmgcs2 in NRK-52E cells seeded on Mfap2-silenced matrix scaffold under CoCl_2_-induced hypoxia. (G) Diagram summarizing conditioned medium (CM) experiments. (H) Western blot assay showing increased CCP3 in NRK-52E cells incubated with Mfap2-deficient CM under Staurosporine stress. (I) Western blot analysis of NRK-52E cells incubated with Mfap2-deficient CM under CoCl₂-induced hypoxia revealed elevated Ngal, Bax, Map2k4, Lats1, and decreased Esr2 and Hmgcs2. (J, K) Under hypoxia, Erb-041 treatment suppressed Ngal expression in NRK-52E cells seeded on Mfap2-silenced matrix scaffold (J) or treated with Mfap2-deficient CM (K). (L) Western blot assay showing reduced CCP3 in NRK-52E cells seeded on an Mfap2-silenced matrix scaffolds following ERb-041 administration. (M) Western blot showing that decreased substrate stiffness (kilopascal, kPa) upregulated Map2k4 and Map4k4 in NRK-52E cells. (N) Conceptual model illustrating how Mfap2 regulates tubular ketogenesis and cell fate through cell–matrix communications, ultimately restoring energy homeostasis and shaping AKI outcomes. TUNEL, terminal deoxynucleotidyl transferase dUTP nick end labeling; Ngal, neutrophil gelatinase-associated lipocalin; Map2k4, mitogen-activated protein kinase kinase 4; Map4k4, mitogen-activated protein kinase kinase kinase kinase 4; Lats1, large tumor suppressor kinase 1; Hmgcs2, 3-hydroxy-3-methylglutaryl-CoA synthase 2.

To further validate these findings, we employed a separate approach to using conditioned medium (CM). NRK-49F fibroblasts were transfected with Dicer-substrate Mfap2 siRNA for 24 hours, and the resulting supernatant was collected as CM to treat cultured tubular cells (Figure 9G). Tubular cells exposed to Staurosporine and incubated with this CM displayed increased cleaved caspase-3 (Figure 9H) and altered expression of the aforementioned proteins under hypoxic stress (Figure 9I). Impressively, treatment with the Esr2 agonist Erb-041 mitigated tubular injury. It suppressed Ngal expression and caspase-3 cleavage in both scaffold- and CM-treated cells (Figure 9, J-L). Furthermore, expression of Map2k4 and Map4k4 increased in response to mechanical stress and was closely correlated with substrate stiffness (Figure 9M). Taken together, these results indicate that Mfap2 deficiency enhanced kidney stiffness and mechanical stress in AKI, leading to Lats1 activation, which represses Esr2 and diminishes Hmgcs2 induction. This cascade ultimately disrupts metabolic adaptation and compromises tubular repair after AKI (Figure 9N).

## Discussion

In the human body, cells continuously sense and respond to mechanical cues that govern antigen recognition(42), cell fate decision(43), cancer metastasis(44), and tissue homeostasis(5). Our study advances the AKI research paradigm by identifying ECM stiffness as a critical, yet underappreciated, driver of early repair. We demonstrate that biomechanical cues embedded within the ECM reboot tubular metabolism, directing the trajectory of AKI repair. Five major findings support this concept: 1) We profiled a comprehensive proteomic map of the DKS following AKI, offering a foundational resource for the field (Figure 1); 2) Mfap2 was identified as an injury-inducible core matrisome component forming a reparative mechanical niche (Figure 2 and 3); 3) we uncovered a direct mechanistic link between ECM stiffness and ketogenesis-driven tubular energy adaptation (Figure 4); 4) A mechano-transduction cascade was revealed in which Mfap2 activates Lats1, independently modulates Esr2 and Hmgcs2, and dictates injury severity (Figure 5-7), and 5) We demonstrated therapeutic efficacy of Esr2 agonists in mitigating AKI (Figure 8). These findings redefine the ECM not as a passive scaffold but as a dynamic conductor of metabolic and regenerative programs during early AKI.

While ECM stiffness is conventionally not considered beneficial for adaptive repair because it often signals fibrotic remodeling and chronic disease progression(45, 46), our findings introduce a new perspective: transient increases in stiffness may facilitate adaptive repair in AKI. The concept originated from growing evidence that early fibroblast activation is a key event after AKI(2, 8, 9). Within hours, activated fibroblasts or pericytes mobilize to injury site(3), initiating ECM synthesis to reinforce tissue and reshape local mechanical cues that influence neighboring cell behavior. Unlike ECM-rich tissues like cartilage and skin(47), a healthy kidney has relatively low baseline stiffness typically ranging from ∼1.8kPa (newborns) to 4.3kPa (adults)(48–50), with increases often signaling maladaptation after injury. In our collagen-coated silicone gel in vitro experiments, tubular cell proliferation marked by Cyclin B1 and D1 peaked at 2.0 kPa and declined with increasing stiffness levels (Figure 1). This inverse relationship suggests that matrix softening promotes regeneration, whereas persistent stiffening favors fibrosis. Similar mechanical regulations of stem cell fate and regeneration have been reported in liver and lung fibrosis(51).

In this study, our time-resolved proteomics revealed a relatively stable DKS composition at day 0 and 1 after AKI, with marked changes by day 3 (Figure 1), indicating ECM stiffness is dynamic and temporally regulated. (Figure 1). Stiffness rose during the inflammatory phase, potentially limiting regenerative capacity, consistent with the clinical finding that patients with stage 3 AKI exhibit increased kidney stiffness(52). Therefore, incorporating the concept of kidney stiffness into AKI research and characterizing the ECM components that govern early repair is a critical next step. In recent years, advances in tissue engineering have enabled novel biomaterial-based platforms to dissect these dynamics. Among them, our DKS platform (Figures 1 and 9) preserves native 3D architecture, immunomodulatory properties, and a rich array of bioactivity for tissue repair(2, 26), offer compelling advantages over synthetic polymer scaffolds(53). Importantly, unlike fibrotic ECM scaffolds which display broad proteomic shifts(46), only six matrix proteins were significantly altered across three time points following AKI in our study, with Mfap2 being the only consistently upregulated core matrisome component (Figure 2). This positions Mfap2 in a sentinel role in which it modulates the mechanical properties of the injured kidney.

The Mfap family comprises five ECM glycoproteins (Mfap1–5) that supports microfibrillar assembly, elastinogenesis, ECM stability, and signal transduction(54). Among them, Mfap2 binds Fibrillin-1(55), a key structural ECM protein mutated in Marfan syndrome(56, 57), an inherited disorder that affects connective tissues in the human body. While previously linked to cancer(58), we reveal Mfap2 as a protective ECM signal that preserves epithelial integrity and facilitates metabolic adaptation in early AKI. Mfap2-deficient mice showed worsened renal outcomes across three AKI models, including impaired function and elevated tubular injury, inflammation, and cell death (Figure 3). Proteomics indicated Mfap2 deficiency selectively suppressed ketogenesis over fatty acid oxidation (FAO), particularly reduced tubular Hmgcs2 (Figure 4). During AKI, as the epicenter of damage, tubules experience mechanical stress as tissue stiffness rises. Its regeneration requires dynamic energy metabolic adaptation to the hypoxic microenvironment, often via hypoxia-inducible factor 1α (HIF-1α) activation(59). As oxygenation improves during recovery, tubules must “reboot” their metabolism toward a more energy-efficient state. While proximal tubules normally rely on FAO for ATP generation(60), during stress, ketone bodies are synthesized from acetyl-CoA by hepatocytes or kidney local stressed cells(32, 61), but Mfap2 deficiency blunted this metabolic adaptation. Of note, phosphorylation of Hmgcs2 at serine 456 enhances catalytic activity under high ketogenic demand(62), but we did not detect changes at this site, suggesting alternative regulatory mechanisms. Nevertheless, ketone bodies may offer a metabolic bypass to dysfunctional FAO machinery, enhancing mitochondrial efficiency and mitigating injury. Furthermore, fibroblasts also undergo metabolic reprogramming during AKI(63), and ECM stiffness is known to influence mitochondrial dynamics, while our data suggest that metabolic changes from Mfap2 deficiency is largely tubular-specific, leaving the impact on fibroblasts an open question.

Interestingly, Mfap2 does not engage classical mechanosensors, such as integrins in tubules, to transduce ECM stiffness into intracellular metabolic signals (Figure 6). This is possibly due to its localization in fibroblasts and interstitial regions, where it contributes to microfibril architecture and elastic fiber assembly, rather than as a direct ligand for tubular integrins. Meanwhile, Mfap2 loss may trigger compensatory upregulation of fibrillin-1 and other microfibrillar or elastic ECM proteins may preserve ECM-integrin interactions. Instead, Mfap2 likely exerts its effects via activation of mechanotransduction signaling pathways Hippo and MAPK, as supported by our DIA-based phosphoproteomics (Figure 6). Upstream kinases Map2k4 and Map4k4 were altered, positioning MAPKs as potential regulators of Lats1, a core kinase of the Hippo pathway. Although canonical Hippo targets Yap/Taz have no changes even if Mfap2 is absent because they remain cytoplasmic and inactive on soft matrices, Lats1 activity itself appears critical in shaping metabolic status and mechanical microenvironment after AKI. Moreover, Map2k4 and Map4k4 actually act as key kinases fueling the JNK and p38 signaling cascades, which suppress the machinery responsible for F-actin polymerization and stabilization. Consistent with these findings, our spatial transcriptomic analyses showed reduced cell-cell communication, distinct gene signatures between fibroblast-neighbor and non-neighbor regions, and dysregulated MAPK upstream factors in Mfap2 KO kidneys compared to WT (Figure 7 and Supplementary Figure S6), further suggesting that this signaling shift promotes cytoskeletal instability, disrupts cell-cell adhesion, and deranges tissue stiffness.

Aligning with prior findings(32, 34), after AKI, repressed Hmgcs2 in the Mfap2-deficiency model is due to downregulated Pparα transcription and sirt5-mediated upregulation of protein succinylation which are commonly regulated by Esr2 (Figure 5), which is validated and has therapeutic potential (Figure 7). A key mechanistic finding is that we identified Lats1 regulates Esr2 but not Esr1 transcription through a Yap/Taz independent manner (Figure 6). A previous study has reported that N-terminal domain of Lats1 can act as an adaptor in structure to form a complex containing Lats1, Esr1, and ubiquitin-ligase substrate receptor Ddb1–cullin4-associated-factor 1 by which Esr1 is degraded to control breast cell fate(35). However, although Esr2 and Esr1 share functional and sequence similarities, their ligand-binding domains and other structural regions exhibit key differences(64), which results in minor changes in Esr2 ubiquitination after knockdown of Lats1 in our current model (Figure 6). Conversely, knockdown of Lats1 substantially amplified Esr2 gene expression in tubular cells. However, Lats1 is a kinase and has no DNA-binding domain, so this regulation might not be via canonical DNA-binding or transcription factor activity. Given it is Yap/Taz independent, it may be through phosphorylating other transcription factors like P53 or Foxo3a or other co-activators or epigenetic modifications, but further exploration and validation are needed to address this limitation.

Our findings unlock new translational possibilities. In AKI patients, shear wave elastography captures elevated kidney stiffness(52), pointing to early Mfap2 induction as a promising matrix remodeling biomarker. When integrated with Ngal and Kim-1 via machine learning algorithms, Mfap2 may improve our ability to accurately predict AKI. From a therapeutic perspective, in addition to Esr2 agonists and exogenous ketone bodies like β-OHB, restoring Mfap2 activity or mimicking its effects using engineered porous scaffolds that replicate the structural and biochemical features of the kidney may enhance tubular cell regeneration and prevent metabolic collapse upon AKI. Yet, translations remain need to overcome key hurdles, including preserving scaffold bioactivity, ensuring effective recellularization, and refining manufacturing processes which should be a shared frontier across regenerative strategies for kidney, lung, liver, bone, nerve, and skin(65).

In summary, this study establishes ECM stiffness as a key early switch in AKI repair, initiating a matrix-triggered metabolic reboot that tips the balance toward regeneration or fibrosis. As AKI disrupts tissue architecture and ECM integrity, effective repair requires not only tubular regeneration but also the restoration of mechanical homeostasis. These insights highlight the imperative to integrate biomechanical and metabolic dimensions into next-generation AKI therapies.

## Methods

The detailed information of methodology, usage of chemical and biological reagents, antibodies, and nucleotide sequences of the primers is presented in the Supplementary Methods.

### Sex as a biological variable

Our AKI study was conducted exclusively in male mice, as female mice exhibit greater resistance to renal ischemia-reperfusion injury and cisplatin-induced toxicity. Human kidney biopsy samples were collected from both male and female subjects. Whether the observations in male mice are applicable to females remains uncertain.

### Statistics

Data are presented as mean ± SEM unless otherwise noted in the figure legends. Statistical analyses were performed using GraphPad Prism 9 (GraphPad Software, San Diego, CA). Two-group comparisons were conducted using a two-tailed Student’s t-test or the Rank Sum Test when data did not meet normality assumptions. For comparisons among multiple groups, one-way ANOVA followed by the Student-Newman-Keuls post hoc test was used. Results are shown as dot plots, with each dot representing an individual data point. A p-value < 0.05 was considered statistically significant.

### Study approval

All animal experiments were performed in accordance with institutional and federal guidelines and approved by the Institutional Animal Care Committee of the University of Connecticut, School of Medicine (Protocol number: AP-200945-0626).

## Data availability

All supporting data are available within the Supporting Data Values XLS file. The raw mass spectrometry (MS) data have been deposited to the ProteomeXchange with identifier PXD064267 and the matrix scaffold MS raw files are available via the MassIVE repository (MSV000087197) at https://massive.ucsd.edu/. The 10X Visium spatial transcriptomics data was submitted to the GEO database with accession GSE299736. All the raw sequencing, imaging and CellRanger filtered matrix files can be downloaded from https://www.ncbi.nlm.nih.gov/geo/query/acc.cgi?acc=GSE299736.

## Authors Contributions

YG and DZ conceived the project. YG and DZ wrote and revised the manuscript. YG, YYW, KZ, JL, HS, CJ, and SM performed most of the in vivo, ex vivo, in vitro studies involving mRNA expression analysis, western blotting assay, immunostaining and imaging. WL, YY, and YL performed proteomic analysis. JJL and SL performed public data mining and spatial transcriptomics analysis. YG, SL, YL, and DZ edited the manuscript. DZ supervised the entire project.

## Acknowledgments

This work was supported by the National Institutes of Health (NIH) grants DK116816, DK128529, DK132059. YG is supported by American Heart Association-Career Development Award (25CDA1435759) and the University of Connecticut InCHIP Faculty Seed Grant. This study was also supported in part by the University of Pittsburgh Center for Research Computing (S10OD028483) and the Jackson Laboratory Cancer Center (P30 CA034196) through the resources provided. We would like to acknowledge Bernard L. Cook III, a Science Editor & Illustrator at UConn Health, for proofreading this manuscript and assisting with illustrations.

