## Supplemental Materials for "Stiffness sensing fuels matrix-driven metabolic reboot for kidney repair and regeneration"

### Supplementary Materials

### Materials and Methods

#### Mice and genotyping

Heterozygous *Mfap2* mice (Product name: S-KO-03113, Strain name: C57BL/6JCya-Mfap2em1) were obtained from Cyagen (Santa Clara, CA). Male and female *Mfap2* heterozygous mice were mated to generate *Mfap2*-wild type (WT) and *Mfap2*-knockout (KO) offspring. Genotyping was performed on tail DNA using standard PCR with the following primers: 5'-TCACCAAGACCACACTCTTGTTA-3', 5'- GGAGCTGGTGTGAGATTCGAG -3', and 5'-ATTACATAGACCTGTGAGGAGGGAC-3'. All mice were born normally at the expected Mendelian frequencies. At baseline, Mfap2 knockout mice were normal in size and did not display any gross physical or behavioral abnormalities.

To evaluate the efficacy of estrogen receptor 2 (Esr2) agonist (Erβ-041) studies, male Mfap2 knockout mice were - intraperitoneallyinjected with Erβ-041 (HY-14933, MCE) at a dose of 1mg/kg body weight/day, starting 2 days prior to IRI model construction. Detailed experimental design is presented in Figure 8A. Blood and kidneys were collected for further analyses.

**Mouse models of AKI**

AS previously described(1), two mouse AKI models were employed: bilateral ischemia-reperfusion injury (BIRI) and Cisplatin injection. For the BIRI model, in brief, after mice were anesthetized, a midline abdominal incision was made, and bilateral renal pedicles were clipped for 25 or 30 minutes using microaneurysm clamps. The mouse body temperature was maintained between 36°C and 37.5°C using a temperature-controlled heating system during the ischemic period. Mice were euthanized at 1 day after IRI, and serum and kidney tissues were collected for various analyses. The cisplatin-induced AKI model involved a single intraperitoneal injection of cisplatin at 30 mg/kg body weight, and then mice were sacrificed 3 days after injection. We used male mice in this study because female mice are relatively resistant to ischemic injury and cisplatin. All animals were housed in standard conditions (69° to 72° F, 12-hour light/dark cycle) in the university animal facility. The study protocol was approved by the Institutional Animal Care and Use Committee at the University of Connecticut School of Medicine.

**Determination of serum creatinine (Scr) and blood urea nitrogen (BUN)**

Serum was collected from mice 1 day after IRI or 3 days after cisplatin injection. Serum creatinine (Scr) and blood urea nitrogen (BUN) levels were respectively determined using the Creatinine (DICT-500) and QuantiChromTM Urea (DIUR-100) assay kits, according to the protocols specified by the manufacturer (BioAssay Systems, Hayward, CA). The levels of Scr and BUN were expressed as milligrams per 100 ml (dL).

**Preparation of Substrates with Defined Stiffness for Cell Culture**

For experiments assessing the impact of substrate stiffness, NRK-52E cells were cultured on CytoSoft® plates of varying elastic moduli (Sigma-Aldrich, #5190-7EA). To enable cell adhesion, the surfaces were coated with PureCol® Type I collagen (Advanced BioMatrix, #5005-100ML) at a concentration of 100 μg/mL. The collagen solution was first warmed to room temperature, and an appropriate volume was added to each well. The plates were then incubated, covered, at room temperature for 1 hour. After incubation, excess collagen solution was aspirated, and the surfaces were rinsed twice with culture medium. Following the second rinse, a thin layer of medium was left to maintain surface hydration. NRK-52E cells were then seeded onto the prepared CytoSoft® surfaces for subsequent experiments.

**Kidney decellularized extracellular matrix (ECM) scaffolds preparation**

Eight-week-old male C57BL/6J mice were purchased from The Jackson Laboratories (Bar Harbor, ME). The BIRI was performed by employing an established protocol as described previously(1). The mice were sacrificed at Days 0, 1, or 3. Kidney were collected and cut into approximately 3 mm slices. The kidney slices were then subjected to decellularization procedures, treated with 0.02% Trypsin/0.05% EDTA, 3% Triton X-100, and 4% Deoxycholic, until all cells were removed and remain decellularized ECM scaffolds.

**Human Kidney Biopsy Specimens**

Human kidney specimens were obtained from diagnostic kidney biopsies performed at the Presbyterian Hospital of the University of Pittsburgh Medical Center. Nontumor kidney tissue from patients with renal cell carcinoma who underwent nephrectomy was used as controls. All patients in the presented study signed informed consent forms before they underwent kidney biopsy or nephrectomy. The Institutional Review Board approved all studies involving human kidney sections at the University of Pittsburgh and the University of Connecticut School of Medicine.

**ATP Measurement**

ATP content in kidney tissue was measured by using the ATP Colorimetric/Fluorometric Assay Kit (ab83355, Abcam, Cambridge, MA), according to the manufacturer’s instructions. Data were normalized for total protein content.

**β-Hydroxybutyrate (β-OHB) Measurement**

The levels of serum β-OHB were measured by using β-Hydroxybutyrate Assay Kit (MAK41-1KT, Sigma-Aldrich), according to the manufacturer's instructions.

**Histology and Immunohistochemical Staining**

Paraffin-embedded mouse kidney sections (3 μm thickness) were prepared by a routine procedure. The sections were stained with periodic acid–Schiff staining reagents by standard protocol. Immunohistochemical staining was performed according to the established protocol as described previously (2). After incubation with primary antibodies at 4°C overnight, the slides were then stained with HRP-conjugated secondary antibody (Jackson ImmunoResearch Laboratories, West Grove, PA). Non-immune normal IgG was used to replace primary antibodies as a negative control, and no staining was visible. Slides were viewed under an Olympus BX43 microscope equipped with a digital camera (Allentown, PA). The image quantification was independently performed by two experienced technicians. The detailed information of antibodies used is presented in Supplementary Table S1.

**Immunofluorescence staining and confocal microscopy**

Kidney cryosections or cell-coated coverslip were fixed with 3.7% paraformaldehyde for 15 min at room temperature. After blocking with 10% donkey serum for 1 hour, the slides were immunostained with primary antibodies or double stained with lotus tetragonolobus lectin (LTL). These slides were then stained with Cy2- or Cy3-conjugated secondary antibody (Jackson ImmunoResearch Laboratories, West Grove, PA). Slides were viewed under an Olympus BX43 microscope equipped with a digital camera or an Olympus FluoView 1000 confocal microscope. The image quantification was independently performed by two experienced technicians. The detailed information of antibodies used is presented in Supplementary Table S1.

**Detection of apoptotic cells**

Apoptotic cell death was determined by using TUNEL staining with a DeadEnd Fluorometric Apoptosis Detection System (Promega, Madison, WI), as we previously described(2).

**Co-immunoprecipitation**

Co-immunoprecipitation was carried out using an established method. Briefly, kidney tissues or normal rat kidney proximal tubular epithelial cells (NRK-52E) stimulated with $CoCl_2$ were lysed on ice in 1 ml non-denaturing lysis buffer that contained 1% Triton X-100, 0.01 mol/l Tris-HCl (pH 8.0), 0.14 mol/l NaCl, 0.025% NaN3, 1% protease inhibitors cocktail, and 1% phosphatase inhibitors cocktail I and II (Sigma). Kidney tissues or cells lysates were incubated overnight at 4°C with 2 mg of anti-Hmgcs2 (sc-393256, Santa Cruz Biotechnology, Dallas, TX) or anti-Esr2 (sc-390243, Proteintech, Rosemont, IL), followed by precipitation with 100 ml of protein A/G Plus-agarose for 3h at 4°C. The precipitated complexes were separated by SDS–polyacrylamide gel electrophoresis and immunoblotted with specific antibodies against succinyllysine (PTM-401, PTM Biolabs, Chicago, IL) or Hmgcs2 (LS-B11023, LsBio, Seattle, WA) or Ubiquitin (ab134953, Abcam, Cambridge, MA), respectively.

**Western Blot Analysis**

Kidney tissues were lysed with radioimmune precipitation assay (RIPA) buffer containing 1% NP-40, 0.1% SDS, 100 μg/ml PMSF, 1% protease inhibitor cocktail, and 1% phosphatase I and II inhibitor cocktail (Cell Signaling Technology, Danvers, MA) in PBS on ice. The supernatants were collected after centrifugation at 13,000×g at 4°C for 15 min. Protein expression was analyzed by western blot analysis as described previouslyS8. The detailed information of antibodies used is presented in Supplementary Table S1.

**Quantitative Real-Time Reverse Transcription PCR (qRT-PCR)**

Total RNA isolation and qRT-PCR were carried out by procedures described previously(1). Briefly, the first strand cDNA synthesis was carried out using a reverse transcription system kit according to the instructions of the manufacturer (Promega). qRT-PCR was performed on an ABI PRISM 7000 sequence detection system (Applied Biosystems, Foster City, CA). The mRNA levels of various genes were calculated after normalizing with β-actin. Primer sequences used for amplifications are presented in Supplementary Table S2.

**Global Proteomics and Phosphoproteomics**

Kidney tissues and decellularized kidney matrix scaffolds were processed following a proteomics workflow published previously (3-5). In brief, WT and Mfap2 KO mice kidneys were lysed using

SDS buffer (4% SDS, 50mM EDTA, 20mM DTT, 2% Tween 20, 100mM Tris-HCl, pH 8.0) and sonication to ensure complete lysis, the tissue then underwent ultrasonic lysis by sonication (Misonix Sonicator 3000 Ultrasonic Cell Disruptor) at 4°C for 10 minutes (with 5 seconds on/off cycles). The lysed samples were then centrifuged at 20,000 x g for 1 hour to remove insoluble material. About 800 μg of proteins were utilized for the subsequent digestion process. Reduction and alkylation were carried out using 10 mM Dithiothreitol (DTT) for 1 hour at 56°C, followed by 20 mM iodoacetamide (IAA) in darkness for 45 minutes at room temperature. The samples were then diluted with 100 mM NH4HCO3 and digested with trypsin (Promega) at a ratio of 1:20 (w/w) overnight at 37°C. The purification of the digested peptides was performed using a C18 column (MaroSpin Columns, NEST Group INC). Five percent of the peptide was utilized for total proteome analysis, while the remaining 95% of the peptide was employed for phosphopeptide enrichment.

The phosphopeptide enrichment process utilized the High-Select Fe-NTA kit (Thermo Fisher Scientific, A32992) following the manufacturer's instruction(6, 7). Briefly, the resins of the spin column were aliquoted and incubated with 250 μg of total peptides for 30 minutes at room temperature, after which they were transferred into the filter tip (TF-20-L-R-S, Axygen). The supernatant was removed by centrifugation. Subsequently, the resins adsorbed with phosphopeptides underwent three washes with 200 μL of washing buffer (containing 80% acetonitrile and 0.1% trifluoroacetic acid), followed by two washes with 200 μL of water to remove nonspecifically adsorbed peptides. The phosphopeptides were then eluted from the resins twice with 100 μl of elution buffer (containing 50% acetonitrile and 5% NH3•H2O). All centrifugation steps were carried out at 500g for 30 seconds. The eluates were dried using a SpeedVac and stored at −80°C before mass spectrometry (MS) analysis.

The samples were measured by data-independent acquisition (DIA) MS method as described previously(8-10) , on an Orbitrap Fusion Tribrid mass spectrometer (Thermo Scientific) coupled to a nanoelectrospray ion source (NanoFlex, Thermo Scientific) and an EASY-nLC 1200 system (Thermo Scientific, San Jose, CA). A 120-min gradient was used for the data acquisition at the flow rate at 300 nL/min with the column temperature controlled at 60 °C using a column oven (PRSO-V1, Sonation GmbH, Biberach, Germany). The DIA-MS method consisted of one MS1 scan and 33 MS2 scans of variable isolated windows with 1 m/z overlapping between windows. The MS1 scan range was 350 – 1650 m/z and the MS1 resolution was 120,000 at m/z 200. The MS1 full scan AGC target value was set to be 2E6 and the maximum injection time was 100 ms. The MS2 resolution was set to 30,000 at m/z 200 with the MS2 scan range 200 – 1800 m/z and the normalized HCD collision energy was 28%. The MS2 AGC was set to be 1.5E6 and the maximum injection time was 50 ms. The default peptide charge state was set to 2. Both MS1 and MS2 spectra were recorded in profile mode. DIA-MS data analysis was performed using Spectronaut v18(11-13), with directDIA algorithm by searching against the SwissProt downloaded mouse fasta file (date). The oxidation at methionine was set as variable modification, whereas carbamidomethylation at cysteine was set as fixed modification. For the phosphorylation data, phosphorylation (S/T/Y) (PTM score >0.75) were set as variable modification as well. Both peptide and protein FDR cutoffs (Qvalue) were controlled below 1% and the resulting quantitative data matrix were exported from Spectronaut. All the other settings in Spectronaut were kept as Default. In phosphorylation data analysis, the identification results were summarized with the PTM score > 0.75, while the quantitative table was filtered with PTM scores > 0.01 to minimize missing values due to phosphosite location for downstream analysis.

**Visium CytAssist spatial transcriptomics**

WT and Mfap2 KO mice ischemic kidneys were quickly collected and embedded in OCT. Then, immersing the OCT-embedded block in isopentane precooled with liquid nitrogen. Four 25um sections from each tissue block were used for total RNA extraction (Qiagen), and the percentage of RNA fragments larger than 200bp as determined by Agilent TapeStation 4200 High Sensitivity DNA ScreenTape (DV200 score) was used as a measure of RNA quality. Tissue blocks with DV200 scores above 30% were used for downstream processing; selected tissues had DV200 scores above 97%. Briefly, 10um sections from each tissue block were placed on microscopy slides (ColorFrost Plus, Fisher) fixed with MeOH, H&E stained, then imaged in brightfield using a NanoZoomer SQ (Hamamatsu) slide scanner at 40x equivalent magnification (Protocol CG000614, 10x Genomics). Each slide was incubated with mouse-specific probe sets provided by the manufacturer for subsequent mRNA labeling, probe transfer using the CytAssist (10x Genomics) onto one capture area of a 6.5mm x 6.5mm Visium CytAssist Slide, and subsequent library generation per the manufacturer's protocol (10x Genomics, CG000495, Rev C). Library concentration was quantified using a Tapestation High Sensitivity DNA ScreenTape (Agilent) and fluorometry (Thermofisher Qubit) and verified via KAPA qPCR. Libraries were pooled for sequencing on an Illumina NovaSeq X Plus 100 cycle 10B flow cell using a 28-10-10-90 read configuration, targeting 100,000 read pairs per spot covered by tissue (approximately 300M per tissue).

**Spatial Transcriptomics: data pre-processing and downstream analysis**

Illumina base call files for all libraries were converted to FASTQs using bcl2fastq v2.20.0.422 (Illumina). For each tissue section and corresponding library, the whole slide brightfield image and

CytAssist image were aligned manually using the Loupe Browser (v6.4.1) via landmark registration. Each whole slide image was uploaded to a local OMERO server where a rectangular region of interest (ROI) containing just the tissue was drawn via OMERO.web and OMETIFF images of each ROI were programmatically generated using the OMERO Python API. FASTQ files, the image registration JSON file, and associated OMETIFF corresponding to high resolution bright field image were used for further processing, including alignment to the GRCm38-specific filtered probe set (10x Genomics Mouse Probeset v1.0.0) using the version 2.1.0 Space Ranger count pipeline (10x Genomics).

The processed spatial transcriptomics data was imported to R/Bioconductor package Seurat for downstream analysis(14). The WT and the Mfap2 KO libraries were first normalized individually and then integrated using the SCTransform function. Following this, the spatial transcriptomics data were clustered based on the expression profile to identify three regional areas. Gene expressions were visualized using DotPlot, SpatialFeaturePlot and rug plot. Subsequently, spatial transcriptomics data were further integrated with the public single-cell RNA-seq data downloaded from GEO database with accession GSE180420(15). Spatial deconvolution was performed and visualized by tool CARD to spatially infer the proportions of different kidney cell types per spot. Based on the cell annotation, tubular cells that are adjacent to (Knn, K-nearest neighbor) and distant from (non-Knn) fibroblasts were identified and compared to identify differentially expressed genes (DEGs). Gene Oncology (GO) pathway analysis was employed on these DEGs to detect enriched pathways associated with Mfap2 knockout.

**Fibroblast decellularized ECM scaffold preparation**

Serum-starved NRK-49F cells under CoCl2 induced-hypoxia stress were transfected with Dicer-substrate Mfap2-siRNA for 24 h. Cells were then treated with EGTA (#3889; Sigma, St Louis, MO) (0.5 mM, PH=7.4) in calcium-free PBS, followed by shaking at 4°C for 1h. The treatment was repeated 3-4 times until all cells were lifted from their underlying matrix. The fibroblast decellularized ECM scaffold was rinsed with PBS and then stored at 4°C for further experiments, as previously described(1, 16). Some ECM scaffolds were collected by scraping with a rubber policeman in loading buffer for western blot assay to detect Mfap2 content. All ex vivo experiments were repeated three times at least.

**Cell culture and treatment**

Normal rat kidney fibroblasts (NRK-49F) and normal rat renal proximal tubular cells (NRK-52E) were obtained from the American Type Culture Collection (ATCC, Manassas, VA). Cells were maintained as described previously (1, 17). For conditioned media (CM) collections, NRK-49F cells were transfected with Dicer-substrate Mfap2 siRNA (rn.Ri. MFAP2.13) for 24 hours and then cultured with serum-free media for 24 hours. The cultured medium was harvested and centrifuged (3000 rpm for 10 min at 4°C). The supernatant was aliquoted and stored at -80°C for subsequent experiments. Serum-starved NRK-52E cells were then treated with CM under Staurosporine (STS, S4400) or $CoCl_2$ (232696; Sigma, St Louis, MO) stress. In addition, NRK-52E cell were transfected with Dicer-substrate Lats1 siRNA or treated with Erβ-041 or Actinomycin D for specific assays. All in vitro experiments were performed three times at least.

**Transmission electron microscopy (TEM)**

Dissected kidney matrix scaffold samples were fixed immediately in 2.5% glutaraldehyde and 2% paraformaldehyde in 0.1 M cacodylate buffer (pH 7.4) for overnight at 4°C. Several 1 mm3 cubes were obtained, washed 5x in cacodylate buffer then post-fixed in aqueous 1% $OsO_4$, 0.8% Ferricyanide for 1 hour. Following 6X $ddH_2O$ washes, the pellet was dehydrated through a graded series of 50%, 75%, 95%, 100% ethanol , and 100% propylene oxide, then infiltrated in 1:1 mixture of propylene oxide: Polybed 812 epoxy resin (Polysciences, Warrington, PA) for 1 hr. After several changes of 100% resin over 24 hours, the pellet was embedded in molds, cured at 37°C overnight, followed by additional hardening at 65°C for two more days. Ultrathin (60 nm) sections were cut on a Leica EM UC7 ultramicrotome, collected on 200 mesh copper grids, stained with 4% uranyl acetate for 10 minutes, followed by 1% lead citrate for 7 min using the Leica EM AC20 automatic stainer. Sections were imaged using a JEOL JEM 1011 transmission electron microscope (Peabody, MA) at 80 kV fitted with a side mount AMT 2k digital camera (Advanced Microscopy Techniques, Danvers, MA). The measurement of foot process width was performed under 15,000x magnification.

**Public data mining**

Public database Tabula Sapiens, a multiple-organ, single-cell transcriptomic atlas of humans, was mined in the study (2). Per organ, single cell gene count matrix was downloaded and loaded into R package Seurat for further analysis (18). Mfap2 expression ratio was calculated by the number of cells with Mfap2 expression over the total number of cells. Other public single-cell nucleus RNA-seq studies on mice were explored in this project (15, 19). The gene-by-cell count matrix per library was downloaded from the GEO database with accession ID GSE180420 and normalized by the *SCTransform* function using the R Seurat package. Then the gene expression across multiple time points was visualized using the DotPlot function.

### Statistics

All data were expressed as mean +/- SEM if not specified otherwise in the legends. Statistical analysis of the data was performed using GraphPad Prism 9 (GraphPad Software, San Diego, CA). Comparison between two groups was made using a two-tailed Student's t-test or the Rank Sum Test if data failed a normality test. Statistical significance for multiple groups was assessed by one-way or two-way ANOVA, followed by the Student-Newman-Keuls test. Results are presented in dot plots, with dots denoting individual values. $P < 0.05$ was considered statistically significant.

**Supplementary Table S1. The information of the applied primary and secondary antibodies**

| Name | Vendor | Category Number | Application |
|---|---|---|---|
| Ngal | Abcam, Cambridge, MA | ab63929 | WB |
| FasL | Santa Cruz Biotechnology, Dallas, Texas | sc-19681 | WB |
| Bax | Santa Cruz Biotechnology, Dallas, Texas | sc-7480 | WB |
| Bax | Cell signaling Technology, Danvers, MA | #14796 | WB |
| Bad | Cell signaling Technology, Danvers, MA | #9268 | WB |
| P-MLKL | Abcam, Cambridge, MA | ab196436 | WB |
| MLKL | Cell signaling Technology, Danvers, MA | #37705 | WB |
| GPX4 | Abcam, Cambridge, MA | ab125066 | WB |
| Cleaved Caspase-3 | Cell signaling Technology, Danvers, MA | #9664 | WB |
| Caspase3 | Cell signaling Technology, Danvers, MA | #9662 | WB |
| Cyclin B1 | Abcam, Cambridge, MA | ab228528 | WB |
| Cyclin D1 | Abcam, Cambridge, MA | ab228528 | WB |
| Cdk2 | Abcam, Cambridge, MA | ab228528 | WB |
| Cdk6 | Abcam, Cambridge, MA | ab228528 | WB |
| PDGFR-β | Cell signaling Technology, Danvers, MA | #3169 | WB |
| Vimentin | Cell signaling Technology, Danvers, MA | #5741 | WB |
| α-SMA | Abcam, Cambridge, MA | ab5694 | WB |
| MFAP2 | Santa Cruz Biotechnology, Dallas, Texas | Sc-166075 | WB |
| MFAP2 | Abcam, Cambridge, MA | AB231627 | IF |
| MFAP2 | MyBioSource, San Diego, LA | MBS8291543 | IHC |
| Hmgcs2 | LsBio, Seattle, WA | LS-B11023 | WB, IHC |
| PPARα | Proteintech Group, Rosemont, IL | 15540-1-AP | WB |
| Sirt5 | Proteintech Group, Rosemont, IL | 15122-1-AP | WB |
| Succinyllysine | PTM Biolabs, Chicago, IL | PTM-401 | WB |
| Hmgcs2 | Santa Cruz Biotechnology, Dallas, Texas | Sc-393256 | Co-IP |
| ESR1 | Proteintech Group, Rosemont, IL | 21244-1-AP | WB |
| ESR2 | Proteintech Group, Rosemont, IL | 14007-1-AP | WB |
| Mapk2K4 | Proteintech Group, Rosemont, IL | 17340-1-AP | WB |
| Mapk4K4 | Proteintech Group, Rosemont, IL | 55247-1-AP | WB |
| Mapk3K3 | Proteintech Group, Rosemont, IL | 21072-1-AP | WB |
| Mapk3K20 | Proteintech Group, Rosemont, IL | 28761-1-AP | WB |
| MapK4K3 | Proteintech Group, Rosemont, IL | 14702-1-AP | WB |
| Rac1 | Proteintech Group, Rosemont, IL | 24072-1-AP | WB |
| Rac2 | Proteintech Group, Rosemont, IL | 0735-1-AP | WB |
| Cdc42 | Proteintech Group, Rosemont, IL | 10155-1-AP | WB |
| RhoA | Proteintech Group, Rosemont, IL | 10749-1-AP | WB |
| Itgα3 | Proteintech Group, Rosemont, IL | 21992-1-AP | WB |
| Itgβ1 | Proteintech Group, Rosemont, IL | 26918-1-AP | WB |
| Itgα6 | Proteintech Group, Rosemont, IL | 27189-1-AP | WB |
| Itgβ3 | Proteintech Group, Rosemont, IL | 18309-1-AP | WB |
| Lats1 | Santa Cruz Biotechnology, Dallas, Texas | Sc-398560 | WB |
| Lats1 | Proteintech Group, Rosemont, IL | 17049-1-AP | WB |
| Lats2 | Proteintech Group, Rosemont, IL | 20276-1-AP | WB |
| Ubiquitin | Abcam, Cambridge, MA | ab134953 | WB |
| Yap | Cell signaling Technology, Danvers, MA | #14074 | WB |
| Taz | Proteintech Group, Rosemont, IL | 23306-1-AP | WB |
| TEAD1 | Cell signaling Technology, Danvers, MA | #12292 | WB |
| Fibronectin | Sigma, St. Louis, MO | F3648 | IF |

| | | | |
|---|---|---|---|
| CD45 | Cell signaling Technology, Danvers, MA | #70257 | IHC |
| CD68 | Abcam, Cambridge, MA | ab283654 | IHC |
| α-Tubulin | Sigma, St. Louis, MO | T9026 | WB |
| GAPDH | Santa Cruz Biotechnology, Dallas, Texas | sc-32233 | WB |
| β-Actin | Santa Cruz Biotechnology, Dallas, Texas | sc-47778 | WB |
| Anti-Mouse IgG | Abcam, Cambridge, MA | ab6789 | WB |
| Anti-Rabbit IgG | Abcam, Cambridge, MA | ab6721 | WB |
| Cy3 Anti-Rabbit | Jackson ImmunoResearch | 711-165-152 | IF |
| Alexa Fluor® 488 Anti-Rabbit | Jackson ImmunoResearch | 711-545-152 | IF |
| Biotin-Anti-Rabbit | Jackson ImmunoResearch | 711-065-152 | IHC |
| F-Actin | Invitrogen, Waltham, MA | R37110 | IF |

WB, western blot; IHC, Immunohistochemical staining; IF, Immunofluorescence staining; Co-IP, Co-immunoprecipitation

**Supplementary Table S2. Nucleotide sequences of the primers used for qRT-PCR**

| Mouse gene | Primer Sequence 5' to 3' | |
|---|---|---|
| | Forward | Reverse |
| IL-18 (M) | GACAGCCTGTGTTCGAGGATATG | TGTTCTTACAGGAGAGGGTAGAC |
| Rantes (M) | GCTGCTTTGCCTACCTCTCC | TCGAGTGACAAACACGACTGC |
| IL-6 (M) | CTTGGGACTGATGCTGGTG | TCCACGATTTCCCAGAGAAC |
| MCP1 (M) | TTAAAAACCTGGATCGGAACCAA | GCATTAGCTTCAGATTTACGGGT |
| TNFα (M) | CCCTCACACTCAGATCATCTTCT | CCCTCACACTCAGATCATCTTCT |
| PPARα (M) | ACCACTACGGAGTTCACGCATG | GAATCTTGCAGCTCCGATCACAC |
| Hmgcs2 (M) | AGCTACTGGGATGGTCGCTA | ACGCGTTCTCCATGTGAGTT |
| FOXA2(M) | CGAGCACCATTACGCCTTCAAC | AGTGCATGACCTGTTCGTAGGC |
| FGF21 (M) | ATCAGGGAGGATGGAACAGTGG | AGCTCCATCTGGCTGTTGGCAA |
| Sirt2 (M) | CGAAGGAGTGACACGCTACATG | GGTGGTACTTCTCCAGGTTTGC |
| Sirt3 (M) | GCTACATGCACGGTCTGTCGAA | CAATGTCGGGTTTCACAACGCC |
| Sirt5 (M) | ATCGCAAGGCTGGCACCAAGAA | CTAAAGCTGGGCAGATCGGACT |
| ESR2 (M) | GGTCCTGTGAAGGATGTAAGGC | TAACACTTGCGAAGTCGGCAGG |
| ESR2 (Rat) | AGGATGTACCACCGAATGCCAAGT | TCCAAGTGGGCAAGGAGACAGAAA |
| β-actin (M) | CAGCTGAGAGGGAAATCGTG | CGTTGCCAATAGTGATGACC |
| β-actin (Rat) | TTCCTGGGTATGGAATCCTG | CTTCTGCATCCTGTCAGCAA |

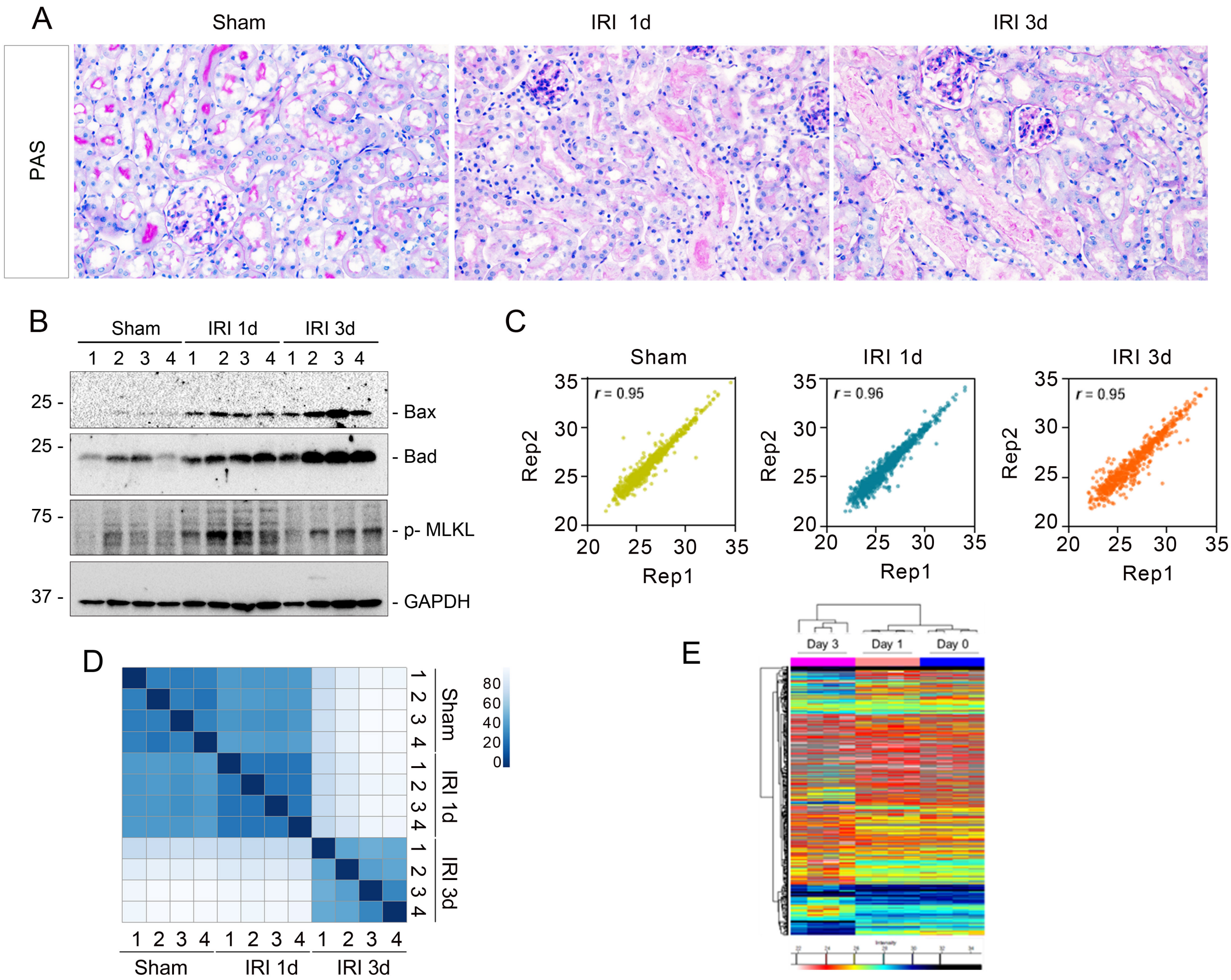


**Supplementary Figure S1: Proteomic profiling of the decellularized kidney matrix (DKS) after ischemic AKI.** (A) Periodic Acid–Schiff (PAS) staining showing morphological alterations in the kidneys after ischemic AKI at day 1 and day 3, compared to sham controls.(B) Western blot analysis showing Bax, Bad, and phosphor-mixed lineage kinase domain-like protein expression in sham and ischemic kidneys after AKI. Numbers indicate individual animals in each group.(C, D) The reproducibility among the biological replicates was generally high, averaging over 0.9 of Pearson correlation for each experimental group. (E) An unsupervised hierarchical clustering of the global proteome in decellularized kidney matrix scaffolds.

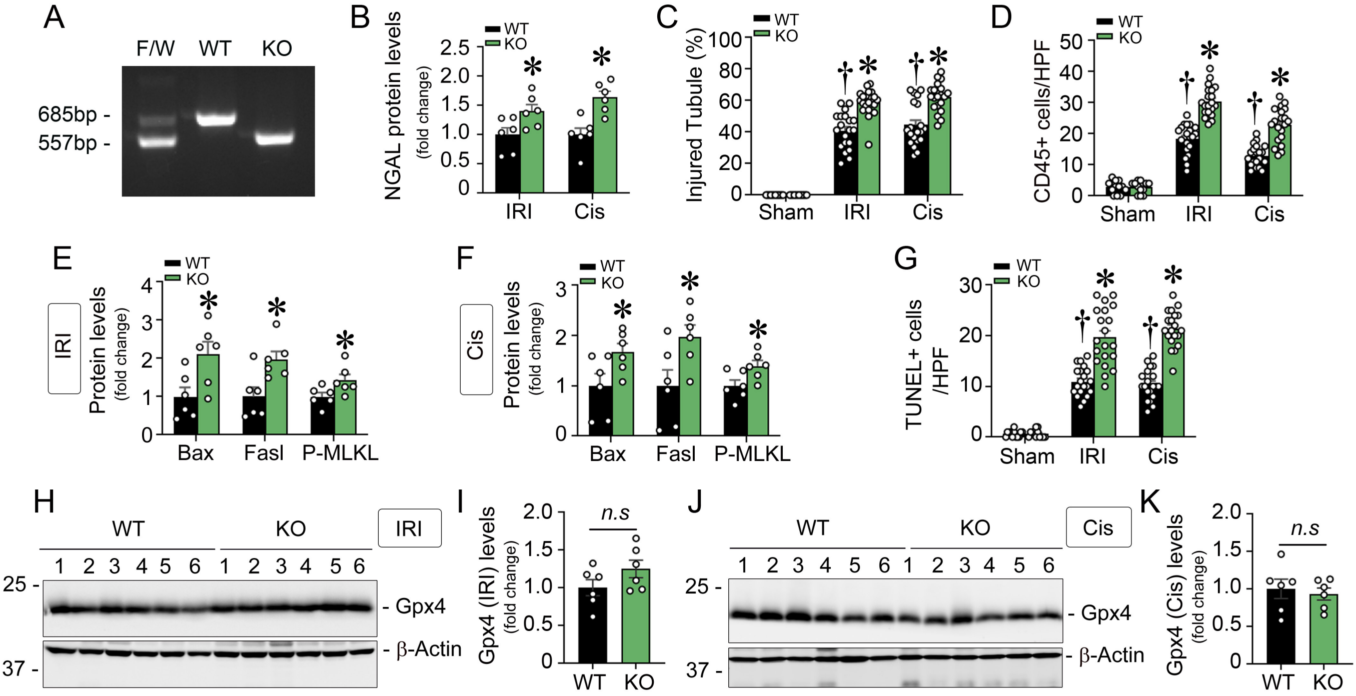


**Supplementary Figure S2: Loss of Mfap2 exacerbates inflammatory cell infiltration and apoptosis but has minimal impact on ferroptosis after AKI**. (A) Genotyping of mice via PCR analysis of genomic DNA. Lane 1: heterozygous control; lane 2: wild type (WT) mice (Mfap2+/+); lane 3: Mfap2 knockout (KO) mice (Mfap2-/-).(B) Quantification of Ngal protein expression in WT and Mfap2 KO mice at day 1 post-IRI or day 3 post-cisplatin injection (* $P < 0.05$; n = 6). (C, D) Quantification of kidney injury scores (C) and CD45+ monocytes infiltration (D) in WT and Mfap2 KO mice at day 1 after IRI or day 3 after cisplatin (Cis) injection. († $P < 0.05$ versus sham, * $P < 0.05$ versus WT; sham, n = 3, IRI or cisplatin, n = 5; four random fields were selected *per* mouse; each dot represents one image). (E, F) Expression levels of Bax, FasL, and pMLKL at day 1 post-IRI (E) or day 3 post-cisplatin injection (F) in WT and Mfap2 KO kidneys (* $P < 0.05$; n = 6). (G) Quantification of TUNEL+ apoptotic cells in WT and Mfap2 KO mice at day 1 post-IRI and day 3 post-cisplatin injection († $P < 0.05$ versus sham, * $P < 0.05$ versus WT; sham, n = 3, IRI or cisplatin, n = 5; four random fields were selected *per* mouse; each dot represents one image). (H-K) Western blot analysis of Gpx4 protein at day 1 post-IRI (H) and day 3 post-cisplatin injection (J), with quantification shown in (I) and (K) (n = 6). Graphs are presented as means ± SEM. Differences among groups were analyzed using unpaired t-tests or one-way ANOVA followed by the Student-Newman-Keuls test. Ngal, neutrophil gelatinase-associated lipocalin; pMLKL, phosphor-mixed lineage kinase domain-like; IRI, ischemia-reperfusion injury; Gpx4, glutathione peroxidase 4; n.s., not significant.

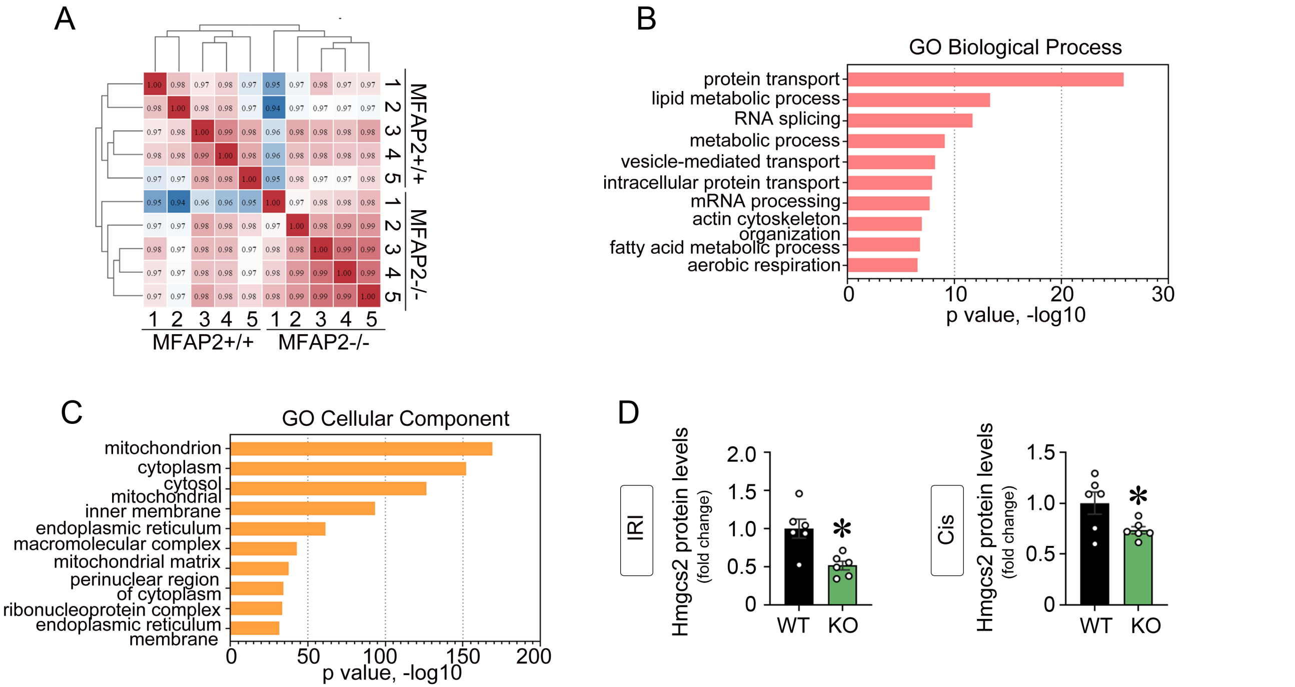


**Supplementary Figure S3: Global proteomic profiling uncovered alternations in the kidney proteome landscape after AKI in Mfap2-deficiency mice.** (A) Correlation of the kidney proteome profiles between WT and Mfap2 KO mice after ischemic AKI. Color scale represents $R^2$ values.(B, C) Gene Ontology (GO) analysis of significantly altered proteins between genotypes, categorized by biological processes (B) and cellular compartments (C); top enriched terms are shown with significance. (D) Quantification of Hmgcs2 protein levels in WT and Mfap2 KO kidneys at day 1 post-IRI (left panel) day 3 post-cisplatin injection (right panel) (* $P < 0.05$; n = 6). Graphs are presented as means ± SEM. Differences among groups were analyzed using unpaired t-tests.

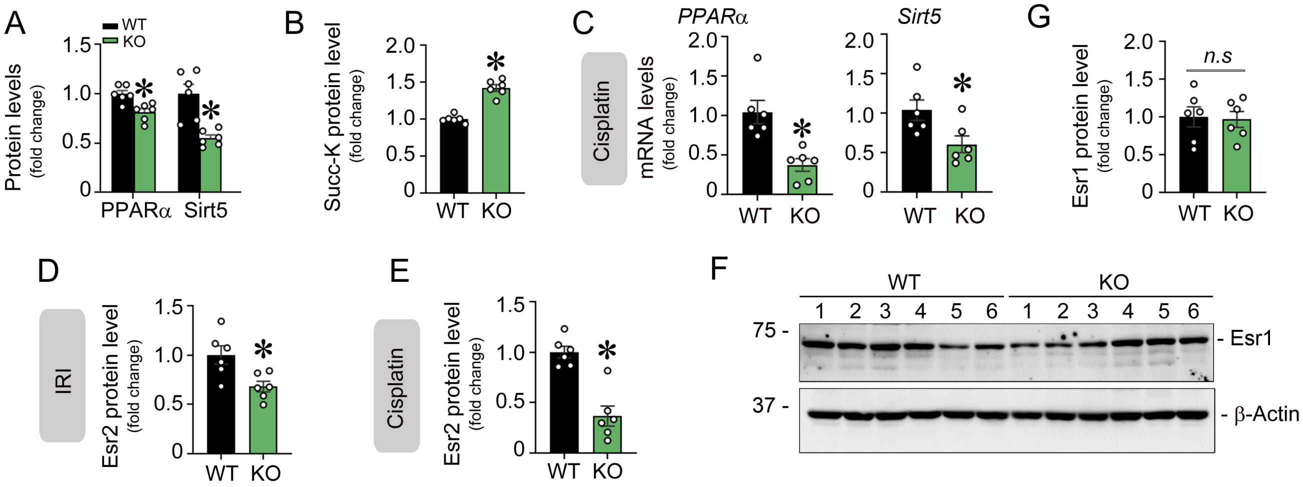


**Supplementary Figure S4: Esr2 regulates Hmgcs2 activity at both transcriptional and posttranslational levels.** (A, B) Quantification of Pparα, Sirt5 protein levels (A) and Succ-K modification (B) in WT and Mfap2 KO kidneys at 1 day after IRI (* $P < 0.05$, n = 6). (C) Quantitative real-time PCR analysis showing *Pparα* and *Sirt5* mRNA expression in WT and Mfap2 KO kidneys at 3 days after cisplatin injection (* $P < 0.05$, n = 6). (D, E) Quantification of Esr2 protein levels in WT and Mfap2 KO kidney at day 1 after IRI (D) or day 3 after cisplatin injection (E) (* $P < 0.05$, n = 6). (F, G) Western blot analysis showing Esr1 protein changes in WT and Mfap2 KO mice at 1 day after IRI (F), with quantitative data shown in (G) (* $P < 0.05$, n = 6). Graphs are presented as means ± SEM. Differences among groups were analyzed using unpaired t-tests. Esr2, estrogen receptor 2; Esr1, estrogen receptor 1; Pparα, peroxisome proliferator-activated receptor α; Succ-K, succinyl-lysine motif; Sirt5, sirtuin 5.

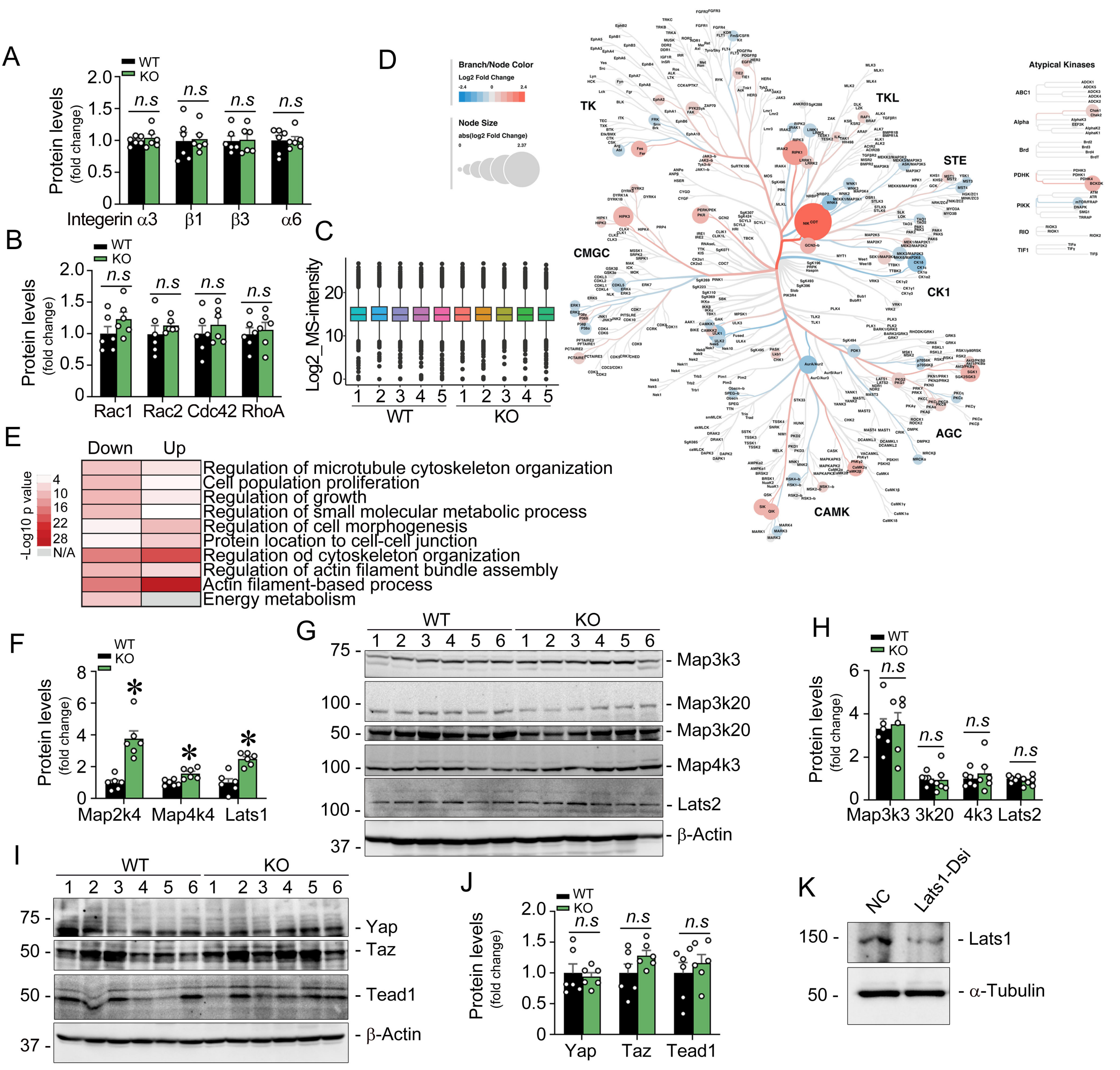


**Supplementary Figure S5: Phosphoproteomic analysis reveals altered signaling pathways upon Mfap2 deletion after IRI.** (A, B) Quantification of integrin subunits (α3, β1, β3, α6) (A) and small Rho GTPases (Rac1, Rac2, CDC42, RhoA) (B) in kidneys from WT and Mfap2 KO mice at day 1 post-IRI (n = 6; n.s., not significant). (C) The distribution of protein intensities in phosphoproteomics. (D) Phylogenetic tree of all protein kinase families detected in the proteomic study.(E) Selected Gene Ontology (GO) pathways enriched in upregulated and downregulated phosphoproteins in ischemic kidneys after Mfap2 deletion. (F) Quantification of Map2k4, Map4k4, and Lats1 protein levels in WT and Mfap2 KO kidneys at day 1 post-IRI (* $P < 0.05$; n = 6). (G, H) Western blot analysis showing expression of Map3k3, Map3k20, Map4k3, and Lats2 in WT and Mfap2 KO kidneys at day 1 post-IRI (G), with quantification shown in (H) (n = 6; n.s., not significant). (I, J) Western blot analysis showing expression levels of Yap, Taz, and Tead1 in WT and Mfap2 KO kidneys at day 1 post-IRI (I), with quantification in (J) (n = 6; n.s., not significant). (K) Western blot analysis demonstrating efficiency of Lats1 knockdown in NRK-52E cells transfected with Dicer-substrate siRNAs (Dsi). Graphs are presented as means ± SEM. Differences among groups were analyzed using unpaired t-tests. Map2k4, mitogen-activated protein kinase kinase 4; Map3k3, mitogen-activated protein kinase kinase kinase 3; Map3k20, mitogen-activated protein kinase kinase kinase 20; Map4k3, mitogen-activated protein kinase kinase kinase kinase 3; Map4k4,mitogen-activated protein kinase kinase kinase kinase 4; Lats1, large tumor suppressor kinase 1; Lats2, large tumor suppressor kinase 2; Yap, yes-associated protein; Taz, transcriptional coactivator with PDZ-binding motif; Tead1, TEA domain transcription factor 1.

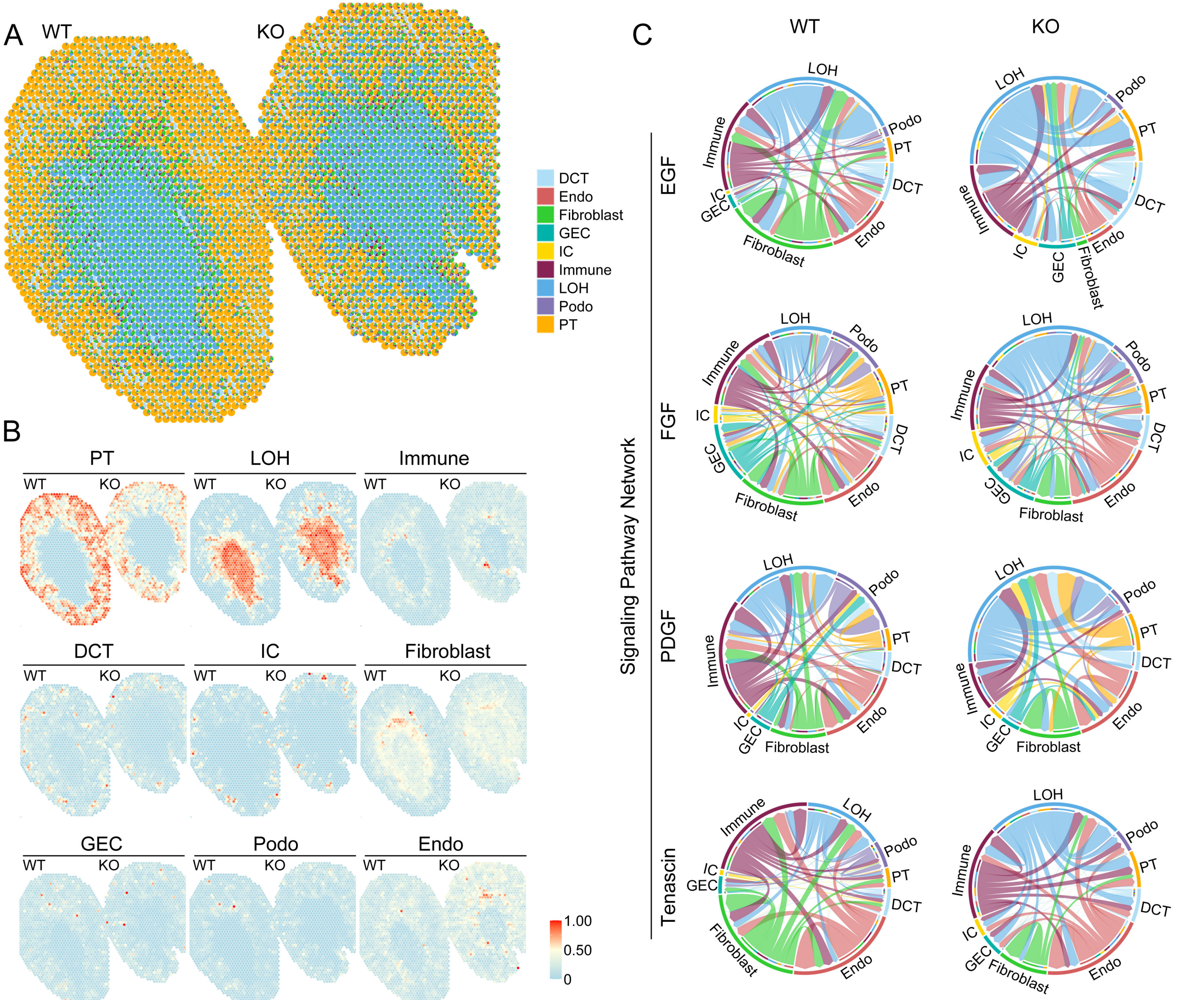


**Supplementary Figure S6: Spatial transcriptomics analysis.** (A) Spatial deconvolution pie graph to show the proportion of cell types at single-cell resolution. DCT, distal convoluted tubule; Endo, endothelial cells; GEC, glomerular endothelial cells; Immune, immune cells; IC, intercalated cells of collecting duct; LOH, loop of Henle; Podo, podocytes; PT, proximal tubular cells. (B) Spatial deconvolution analysis to reveal the ratio of each cell type for all the spots. (C) Spatial network analyses showing alteration of EGF-, FGF-, PDGF-, and Tenascin-mediated signaling connections across nephron and cellular compartments. EGF, epidermal growth factors; FGF, fibroblast growth factor; PDGF, platelet-derived growth factor.

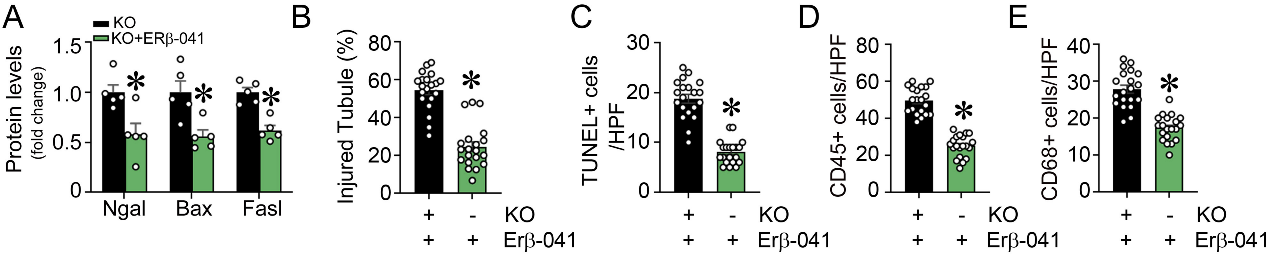


**Supplementary Figure S7: Esr2 agonist attenuates ischemic AKI.** (A) Quantification of protein expression of Ngal, Bax, and FasL in Mfap2 KO kidneys with or without treatment of Esr2 agonist (ERβ-041) at 1 day after IRI. (* *P* < 0.05, n = 5). (B-E) Quantification of kidney morphological injury (B), TUNEL staining (C), CD45 immunostaining (D), and CD68 immunostaining (E) in Mfap2 KO mice treated with or without ERβ-041 at 1 day after IRI. (* *P* < 0.05. Sham, n = 3; IRI, n = 5; 4 random images were selected *per* mouse; each dot represents the injury score of a single image). Graphs are presented as means ± SEM. Differences among groups were analyzed using unpaired t-tests. IRI, ischemia-reperfusion injury; Ngal, neutrophil gelatinase-associated lipocalin; TUNEL, terminal deoxynucleotidyl transferase dUTP nick-end labeling.

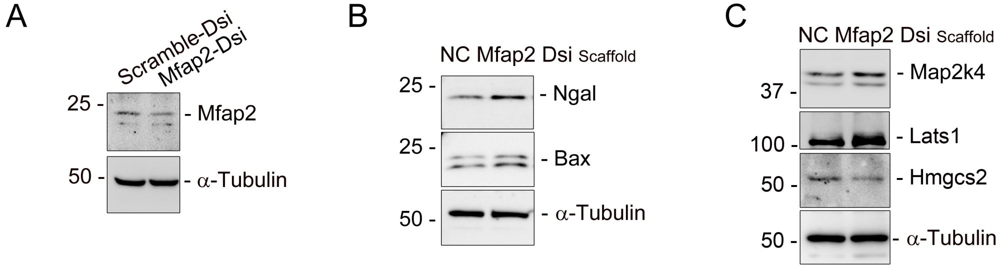


**Supplementary Figure S8: Knockdown of Mfap2 suppresses tubular ketogenesis ex vivo and in vitro.** (A) Western blot analysis confirming Mfap2 knockdown efficiency in normal rat kidney fibroblasts (NRK-49F) transfected with Mfap2 Dicer-substrate siRNAs (Mfap2-Dsi), compared to controls (Scramble-Dsi). (B, C) Western blot analysis of Ngal and Bax (B), as well as Map2k4, Lats1, and the ketogenic enzyme Hmgcs2 (C) in normal rat proximal tubular epithelial cells (NRK-52E) cultured on matrix scaffolds derived from Mfap2-knockdown fibroblasts. Ngal, neutrophil gelatinase-associated lipocalin; Map2k4, mitogen-activated protein kinase kinase 4; Lats1, large tumor suppressor kinase 1; Hmgcs2, 3-hydroxy-3-methylglutaryl-coenzyme A synthase 2.
